# Combining ancestral reconstruction with folding-landscape simulations to engineer heterologous protein expression

**DOI:** 10.1101/2021.07.16.452635

**Authors:** Gloria Gamiz-Arco, Valeria A. Risso, Eric A. Gaucher, Jose A. Gavira, Athi N. Naganathan, Beatriz Ibarra-Molero, Jose M. Sanchez-Ruiz

## Abstract

Obligate symbionts exhibit high evolutionary rates and extensive sequence divergence. Here, we use the thioredoxin from *Candidatus Photodesmus katoptron*, an uncultured symbiont of flashlight fish, to explore evolutionary and engineering aspects of protein folding in heterologous hosts. The symbiont protein is a standard thioredoxin in terms of 3D-structure, stability and redox activity. However, its refolding *in vitro* is very slow and its expression in *E. coli* leads to insoluble protein. By contrast, resurrected Precambrian thioredoxins express efficiently in *E. coli*, plausibly reflecting an ancient adaptation to unassisted folding. We have used a statistical-mechanical model of the folding landscape to guide back-to-ancestor engineering of the symbiont protein. Remarkably, we find that the efficiency of heterologous expression correlates with the *in vitro* refolding rate and that the ancestral expression efficiency can be achieved with only 1-2 back-to-ancestor replacements. These results demonstrate a sequence-engineering approach to rescue inefficient heterologous expression, a major biotechnological bottleneck.

## Introduction

Many proteins of industrial and therapeutic interest are produced in heterologous hosts using recombinant DNA technology^1–3^. Moreover, heterologous expression is unavoidable in metagenomics studies aimed at the functional characterization and biotechnological exploitation of proteins from organisms that cannot be cultured in the lab^4,5^, *i.e.,* the majority of the bacteria in soil^6^. Unfortunately, over-expression of a foreign gene poses a serious challenge to an organism and may lead to non-functional species, such as misfolded protein, proteolyzed protein or, more typically, insoluble protein aggregates^7^. These problems may be alleviated by using engineered hosts that have been modified for instance to minimize protease activity or to over-express molecular chaperones that assist correct folding^2,8^. Despite advances in host engineering, heterologous expression of functional proteins remains a major biotechnological bottleneck. For instance, about half of the proteins targeted in structural genomics initiatives could not be purified^9^.

Ancestral sequence reconstruction (ASR) uses phylogenetic analyses and sequences of modern protein homologs to compute statistically plausible approximations to the corresponding ancestral sequences^10,11^. During the last *∼*25 years, proteins encoded by reconstructed sequences (“resurrected” ancestral proteins) have been widely used as tools to address important problems in molecular evolution^12–16^. They have also been found to provide new possibilities for protein biomedical applications and protein engineering^17–21^. Ancestral proteins may considerably differ from their modern counterparts in terms of sequence and their experimental preparation necessarily involves heterologous expression, as the ancient original hosts are not available. Therefore, the fact that many resurrected ancestral proteins have been purified and studied experimentally emerges as remarkable in itself, even after acknowledging the obvious publication bias in favour of positive experimental outcomes.

Moreover, a substantial number of studies have actually reported improved heterologous expression of ancestral proteins as compared with their modern counterparts. Examples include: phosphate-binding protein^22^, periplasmic binding protein^23^, serum paraoxonase^24^, coagulation factor VIII^25^, titin^26^, haloalkane dehalogenases^27^, cytidine and adenine base editors^28^, diterpene cyclase^29^, rubisco^30^, endoglucanases^31^, L-amino acid oxidases^32^, laccases^33^, front-end Δ6-desaturases^34^ and fatty acid photo-decarboxylases^35^. On a related note, recent studies on ancestral proteins have noted an improved capability to yield crystals suitable for X-ray structural determination^36,37^.

Regardless of the specific mechanisms responsible for efficient ancestral folding in modern organisms, it is clear that ancestral reconstruction may provide a basis for the sequence engineering of efficient heterologous expression. Yet, reconstructed ancestral sequences typically display extensive differences with respect to the corresponding modern sequences while, in many cases, researchers will be interested in minimally modifying the targeted modern sequence, in such a way that the properties of the encoded protein are barely altered. Minimal sequence perturbation would be particularly desirable when targeting recent adaptations, as, for instance, in metagenomics efforts at contaminated sites aimed at obtaining pollutant-degrading enzymes^38^. Overall, it is of interest to determine whether rescue of inefficient heterologous folding can be engineered on the basis of a few selected back-to-ancestor mutations.

Obligate symbionts typically display high evolutionary rates^39,40^ and their proteins may be expected to differ considerably from their modern and ancestral homologs in terms of both sequence and properties. Consequently, the outcome and implications of evolutionary processes may become apparent even in small symbiont proteins that are amenable to detailed biomolecular characterization. Thioredoxins are small proteins (about 110 amino acid residues) that function as general redox catalysts in all known cells^41^. Here we use the thioredoxin from *Candidatus Photodesmus katoptron*, an uncultured symbiont of flashlight fish, as a simple model to explore evolutionary and engineering aspects of protein folding in heterologous hosts.

Flashlight fish (*Anomalopidae*) use light from sub-ocular bioluminescent organs to communicate, hunt prey and disorient predators^42^. Light is produced by luminous bacteria of the *Vibrionaceae* family of *Proteobacteria.* These bacteria are obligate symbionts and have not been cultured in the lab, thus replicating a fundamental scenario of metagenomics studies. Still, their genomes can be sequenced, since the sub-ocular organs of *Anomalopidae* fish harbour large numbers of *Vibrionaceae* in the absence of other bacteria^43^. *Candidatus Photodesmus katoptron*, the luminous bacterium of the *Anolamops katoptron* fish, shows extensive genome reduction and it is highly evolutionary divergent^44–46^. As expected from the high evolutionary rate of its original host, the sequence of the thioredoxin from *Candidatus Photodesmus katoptron* (*CPk* thioredoxin from now on) differs substantially from all known sequences of thioredoxins from other species. Despite this observation, it is similar to other modern thioredoxins (in particular, to its *E. coli* homolog) in terms of 3D-structure, stability and redox activity. However, as described below, its folding outside the original host appears severely impaired.

*CPk* thioredoxin displays a very slow refolding *in* vitro, reaching the native state in the time scale of hours^47^ and its expression in *E. coli* at 37°C leads mostly to insoluble protein. By contrast, resurrected Precambrian thioredoxins have been extensively studied^47–51^ and have been found to fold fast *in vitro* and efficiently in *E. coli* despite their huge sequence differences with modern thioredoxins in general and *E. coli* thioredoxin in particular.

Albeit inefficient, the heterologous folding of *CPk* thioredoxin in *E. coli* is not fully impaired and leads to about 20% of soluble protein at 37°C, a yield that can be increased by carrying out the expression at lower temperatures to decrease protein aggregation^52^. This is a crucial feature that allows us to interrogate the folding properties of *CPk* thioredoxin variants. That is, the effect of sequence modifications on heterologous folding efficiency at 37 °C can be determined and correlated with the biomolecular properties of the corresponding variants of *CPk* thioredoxin, since these variants can actually be prepared in the lab.

We have used computational modelling of the folding landscape to guide back-to-the-ancestor engineering of *CPk* thioredoxin. Specifically, we have used a recently-developed^53^, block version of the Wako-Saitô-Muñoz-Eaton statistical-mechanical model of the folding landscape^54–57^ to determine regions of the symbiont thioredoxin that are likely to be unfolded in aggregation-prone intermediate states^58^ and we have performed back-to-ancestor sequence-engineering targeted to those regions. The ancestral protein we have used as reference is LPBCA thioredoxin, a putative Precambrian thioredoxin that we have previously characterized in detail^47–51^ and that folds efficiently in *E. coli*, despite having only 58% sequence identity with *E. coli* thioredoxin.

Our current study includes several modern/ancestral chimeras, as well as a many single-mutant, back-to-ancestor variants, and allows us to generate several conclusions of general interest:

Folding in the heterologous host is likely akin to unassisted folding. This is supported by (i) the success of the approach used, which involves computational modelling of the unassisted folding landscape, (ii) the fact that the efficiency of heterologous expression correlates with the *in vitro* folding rate, (iii) the very limited rescue of inefficient heterologous expression by chaperone over-expression. Consequently, it appears plausible that ancestral folding efficiency reflects an adaptation to ancient unassisted folding^47^.

Stabilization does improve heterologous folding efficiency, but this cannot be explained by global stabilization alone. Rather, it is linked to specific stabilizing mutations at crucial positions in late-folding regions.

Although the sequences of the ancestral LPBCA thioredoxin and the modern *CPk* thioredoxin differ at 60 positions, the ancestral folding efficiency can be re-enacted in the symbiont thioredoxin with only 1-2 back-to-ancestor mutations. This result provides proof of concept for a minimal-perturbation, sequence-engineering approach to rescue inefficient heterologous folding with potential application in metagenomics.

## Results and discussion

### Sequences and biomolecular properties of the modern and ancestral thioredoxins studied in this work

Our current study utilizes the thioredoxin from the symbiont *Candidatus Photodesmus katoptron* (*CPk* thioredoxin), a modern *E. coli* homolog and a resurrected ancestral thioredoxin corresponding to the last common ancestor of the cyanobacterial, *Deinococcus* and *Thermus* groups, a Precambrian phylogenetic node dated at ∼2.5 billion years ago. This LPBCA thioredoxin, as well as other resurrected Precambrian thioredoxins, has been previously characterized in detail^47–51^.

*CPk* thioredoxin is highly divergent at the sequence level. A BLAST search in the non-redundant protein sequences (nr) database using as query the sequence of *CPk* thioredoxin yields a *vibrio* protein with only 75% identity as the closest hit. Sequence identity of this thioredoxin from *Candidatus Photodesmus katoptron* (belonging to the *Vibrionaceae* family of *Proteobacteria*) with the thioredoxin from *E. coli* (belonging to the *Enterobacteriaceae* family of *Proteobacteria*) is even lower: 69%. The ancestral LPBCA thioredoxin displays even lower sequence identity to both modern proteins: 57% and 45% with the thioredoxins from *E. coli* and *Candidatus Photodesmus katoptron*, respectively.

Despite the extensive sequence differences (Fig. 1a), the three proteins share the thioredoxin fold (Fig. 1b). The 3D-structures of *E. coli* thioredoxin and LPBCA thioredoxin have been previously reported^49,59^. The determination of the X-ray structure for *CPk* thioredoxin has been addressed in this work. However, despite numerous attempts using different crystallization conditions and approaches, we failed to obtain crystals of diffraction quality for the wild-type *CPk* thioredoxin (see Methods for details). The structure shown in Fig. 1b for *CPk* thioredoxin actually corresponds to an engineered version of the protein in which a short loop (70-77) of the symbiont protein has been replaced by the corresponding loop in the ancestral LPBCA thioredoxin (see details below), a replacement which involves only 4 mutational changes. This variant of *CPk* thioredoxin did produce crystals suitable for diffraction and led to structure at 2.85 Å resolution.

**Figure 1.**
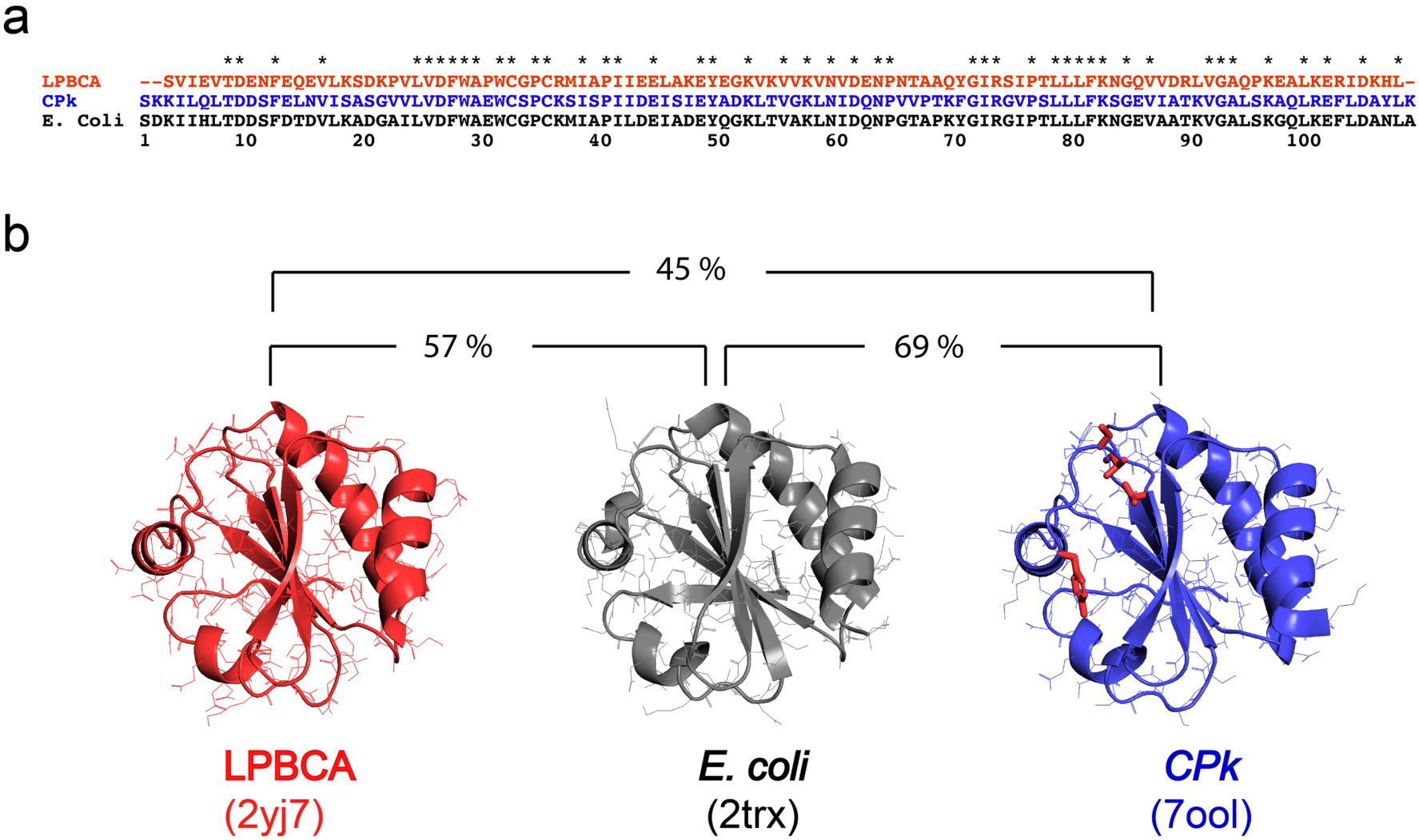
Sequences and structures of modern and ancestral thioredoxins. **a** Alignment of sequences from modern thioredoxins from *E. coli* and *Candidatus Photodesmus katoptron* (*CPk*), and a resurrected ancestral thioredoxin corresponding to the last common ancestor of the cyanobacterial, *Deinococcus* and *Thermus* groups (LPBCA thioredoxin). Positions with identical residues in the three sequences are labelled with asterisks. **b** 3D-structures for the three thioredoxins studied. The experimental structure of *CPk* thioredoxin actually corresponds to a variant with four back-to-ancestor replacements (highlighted in red; see main text for details). The PDB identifiers are shown, as well as the sequence identity percentages between the three proteins.

The agreement between the three structures shown in Fig. 1b is consistent with previous structural work that supported conservation of thioredoxin structure over the span of life on Earth^49^. *In vitro* redox activity, as determined by the insulin aggregation assay and by the assay with thioredoxin reductase coupled to DTNB, is also similar for the three thioredoxins (Fig S1). The two modern thioredoxins display similar high stability as shown by midpoint of about 8M for urea denaturation experiments^47^ and by denaturation temperatures above 80°C (see details below). The ancestral LPBCA thioredoxin is a hyperstable protein that cannot be denatured by urea at room temperature and that has a denaturation temperature of about 123°C^48,60^.

### *In vitro* folding behaviour of the modern and ancestral thioredoxins studied in this work

*In vitro* folding of thioredoxins has been known for many years to be a kinetically complex process involving intermediate states and parallel channels to arrive at the native state^61,62^. Such complexities reflect ruggedness of the folding landscape and are revealed by multi-exponential folding kinetics and rollovers in the folding branches of Chevron plots (*i.e.,* plots of folding-unfolding rate constant versus denaturant concentration). Fig. 2a shows Chevron plots for the three thioredoxins studied here^47^. To identify the kinetic phase that leads to the native state, *i.e.,* the kinetic phase that defines the time-scale of refolding, we used double-jump unfolding assays^47,63,64^, which allow for a direct determination of the amount of native protein during the folding experiments. These assays (see Methods for details) are a specific instance of the well-known “jump assays” that were developed by pioneers of the *in vitro* protein folding field to resolve the kinetic complexities of folding processes^65,66^. While the interpretation of the time dependence of a protein physical property may not be straightforward, double-jump unfolding assays lead to a profile for the fraction at native state *versus* time and reveal immediately the time scale in which the native state is reached upon refolding *in vitro*. Such profiles are given in Fig. 2b and 2c for the three thioredoxins studied. It is clear that folding *in vitro* of *CPk* thioredoxin is substantially slower than the folding of *E. coli* thioredoxin and LPBCA thioredoxin, as discussed above. Furthermore, the *in vitro* folding of *CPk* thioredoxin is also inefficient, as shown by the fact that substantial amounts of protein fail to reach the native state (Fig. 2b). The effect is more pronounced the higher the total concentration of protein, suggesting that *in vitro* folding inefficiency is linked to protein aggregation (which is, in fact, observed visually). The rate constants derived using double-jump unfolding assays (closed symbols in the plots of Fig. 2a) indicate that the native state is reached mostly in the slow kinetic phase detected by fluorescence.

**Figure 2.**
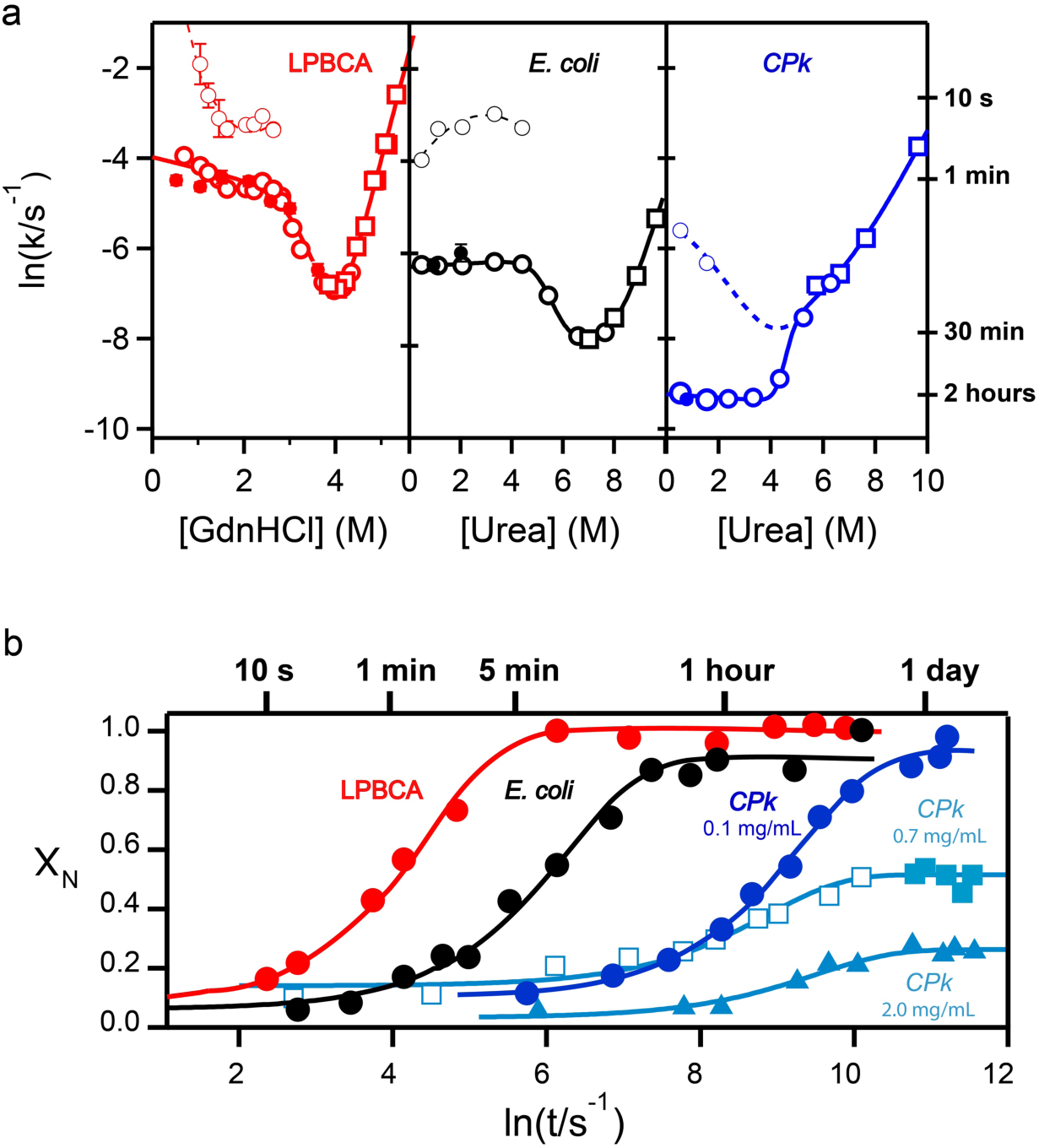
*In vitro* folding of modern and ancestral thioredoxins. **a** Chevron plots of folding-unfolding rate constant at pH 7 and 25°C *versus* denaturant concentration for the three thioredoxins studied. Urea is used as denaturant for *E. coli* and *CPk* thioredoxins. LPBCA thioredoxin, however, is highly stable and cannot be denatured by urea at 25°C. The chevron plot using the stronger denaturant guanidinium hydrochloride is shown for this protein. Still, the folding rates for LPBCA thioredoxin at low denaturant concentration obtained with urea and guanidine are in good agreement^47^. Circles and squares refer to experiments performed in the folding and unfolding directions, respectively. Error bars are standard errors derived from fits to the experimental profiles and are not shown when they are smaller than the size of the data point. Data are taken from Gamiz-Arco et al.^47^ and were derived from fluorescence kinetic profiles. These profiles are often multiphasic in the folding direction, with the slow folding phase leading to the native state, as shown by the agreement with the folding rates (closed symbols) derived using double-jump unfolding assays. Lines shown are meant to guide eye. **b** Profiles of fraction on native state *vs.* time obtained by using double-jump unfolding assays. Experiments were performed at pH 7, 25°C in the presence of 1M urea (see Methods for details). Protein concentration was 0.1 mg/mL, except for the profiles for *CPk* thioredoxin at 0.7 mg/mL and 2 mg/mL. The open symbols in the profile for *CPk* thioredoxin at 0.7 mg/mL are taken from Gamiz-Arco et al.^47^. The lines represent the best fits of a single exponential. Note that an exponential has a sigmoidal shape in a plot versus ln t. In both **a** and **b**, typical values of the half-life time (calculated as the inverse of the first-order rate constant) are indicated to highlight the large differences in folding time-scale between the three proteins.

### Efficiency of the heterologous expression in *E. coli* of modern and ancestral thioredoxins

We determined the efficiency of expression in *E. coli* at 37°C as the ratio of soluble protein to total protein determined after overexpression for 3 hours, as recommended by standard protocols (see Methods for details).

As was to be expected, expression of *E. coli* thioredoxin in *E. coli* is highly efficient, leading to essentially 100% soluble protein. Remarkably, the expression of the ancestral LPBCA thioredoxin is also highly efficient, despite the extensive sequence differences with *E. coli* thioredoxin (only 57% sequence identity). On the other hand, expression of *CPk* thioredoxin in *E. coli* only leads to about 20% soluble protein (Fig. 3).

**Figure 3.**
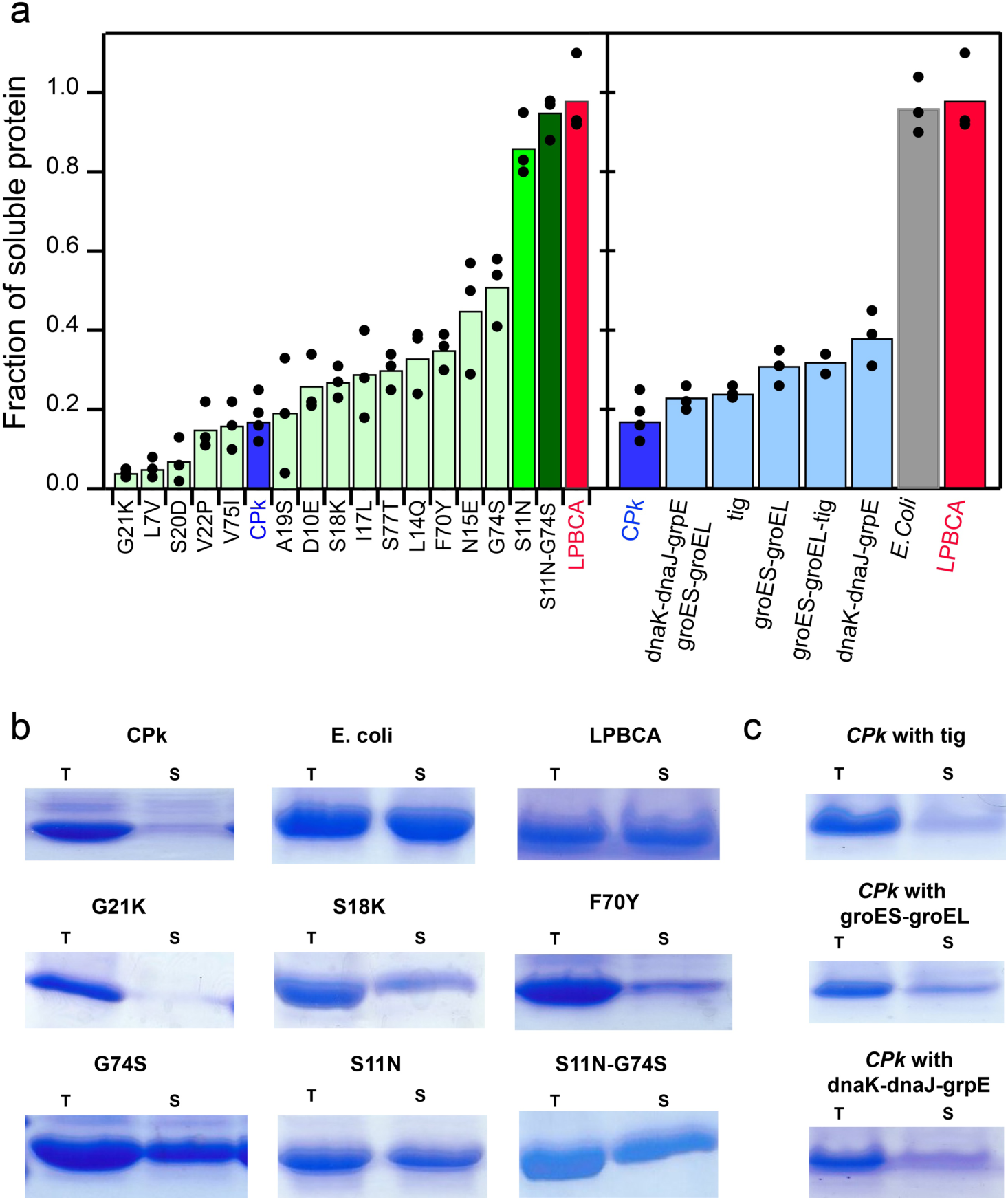
*In vivo* expression of modern and ancestral thioredoxins. **a** Fraction of soluble protein obtained upon expression of thioredoxin proteins in *E. coli* at 37°C. The box at the left displays data for LPBCA thioredoxin, *CPk* thioredoxin, variants of the latter with single mutations and one double-mutant variant (S11N-G74S). The box at the right displays data for *E. coli* thioredoxin, LPBCA thioredoxin, *CPk* thioredoxin and data for the later with over-expression of various chaperone teams. Bars represent the average of several independent determinations and the individual values are also shown. **b** Illustrative examples of the experimental determination of the fraction of soluble protein for LPBCA thioredoxin, *E. coli* thioredoxin, *CPk* thioredoxin and variants of the latter. **c** Representative examples of the experimental determination of the fraction of soluble protein for *CPk* thioredoxin with over-expression of chaperone teams. In both **b** and **c**, T and S represent “total” and “soluble”, respectively. For illustration, only sections of the SDS-PAGE gels with the thioredoxin bands are shown. Complete gels are shown Figures S10-S13.

Chaperone over-expression is a common host-engineering strategy to improve heterologous protein expression^2,8^. We attempted to rescue the inefficient folding of the *CPk* thioredoxin in *E. coli* by complementation with plasmids containing genes of the following *E. coli* chaperones: trigger factor, groES, groEL, dnaK, dnaJ. We used five separate combinations of these chaperones, as shown in Figure 3a. Only very moderate enhancements of *in vivo* folding efficiency were observed (Fig. 3).

### A plausible evolutionary narrative

The experiments described above (Figs. 2 and 3) reveal a striking disparity between the folding behaviour of the *CPk* thioredoxin and LPBCA thioredoxin outside their original hosts. The ancestral protein folds fast *in vitro* and its expression in *E. coli* is efficient. By contrast, expression of the symbiont thioredoxin in *E. coli* produces a substantial amount of insoluble protein and refolding experiments show that it reaches the native state *in vitro* in the time scale of hours. This slow and inefficient folding can hardly be assumed to correspond to the situation *in vivo* in the original host. Note that the synthesis of a *∼*100 residue protein by bacterial ribosomes takes about 5 seconds^58^. Obviously, it is difficult to understand that a small protein, which can in principle fold fast, has been selected during evolution to fold in its original host in a time scale *∼*3 orders of magnitude above the time required for synthesis in the ribosome. A much more likely scenario is that folding of *CPk* thioredoxin in its original host is fast, plausibly allowing for co-translational folding, and efficient. This fast/efficient *in vivo* folding would obviously be the result of the interaction of the protein with the cellular folding assistance machinery, mainly the ribosome itself^67,68^ and the ribosome-binding chaperones (the trigger factor), which would guide and assist co-translational folding, although a role for downstream chaperones and the specific environment in the symbiont (pH, redox, etc.) cannot be ruled out. Such assistance, however, would not be available for *CPk* thioredoxin in the heterologous *E. coli* host, since co-evolution in the original host would lead to its adaptation to the folding assistance machinery of the symbiont. Note that, not only the symbiont thioredoxin, but also most of the symbiont chaperones and ribosomal proteins are highly divergent at the sequence level (see Tables S2 and S3). Therefore, co-evolution of interacting symbiont proteins is a likely scenario.

In view of the above reasoning, the following evolutionary narrative appears plausible. The most ancient proteins emerged before an elaborated folding-assistance machinery was available. Therefore, ancestral folding efficiency relied on fast unassisted folding that limits the transient population of aggregation-prone partially-unfolded states. Consequently, efficient folding of resurrected ancestral proteins in modern organisms may plausibly reflect an ancient adaptation to unassisted folding. Of course, efficient unassisted folding would no longer be a useful feature after the evolutionary emergence of cellular folding-assistance, which would thus allow the evolutionary acceptance of mutations that impair the ancestral feature. Such degradation of unassisted folding would be of no consequence for folding in the original host, but will lead to inefficient expression in a heterologous host, where folding assistance would not be available due to co-evolution in the original host.

Evolutionary narratives are necessarily speculative. Still, the narrative we have proposed above has the merit of immediately suggesting an approach to the sequence-engineering of heterologous expression. That is, suitable back-to-ancestor mutations could lead to a more efficient heterologous expression. Furthermore, since folding in the heterologous host is, at least to some extent, unassisted, computational modelling of the unassisted folding-landscape may be used to guide back-to-ancestor engineering for efficient heterologous expression. Computational modelling of the folding landscape for thioredoxins is described in the next section.

### Computational modelling of the folding landscape for modern and ancestral thioredoxins

Here we use a recently developed version^53^ of the Wako-Saitô-Muñoz-Eaton (WSME) statistical model of protein folding^54–57^ to assess the main features of the folding landscape of modern and ancestral thioredoxins (Fig. 4).

**Figure 4.**
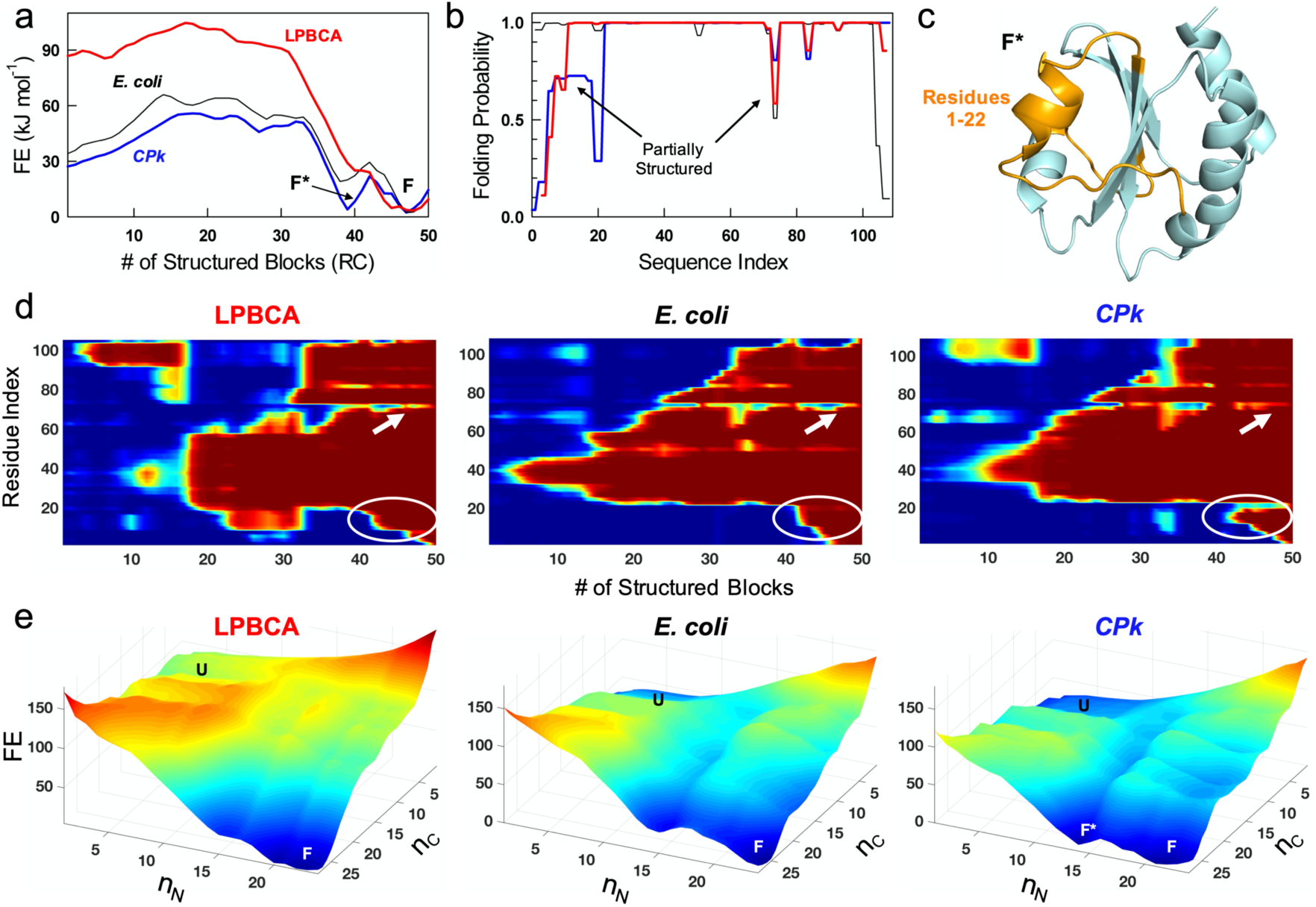
Statistical mechanical modelling of the folding landscape for modern and ancestral thioredoxins. A block version of the Wako-Saitô-Muñoz-Eaton model was used in all the calculations shown here. **a** Profiles of free energy versus number of structured blocks at 37°C for the modern thioredoxins from *E. coli* and *Candidatus Photodesmus katoptron* (*CPk*), and the ancestral LPBCA thioredoxin. Note that a partially-unfolded intermediate (F* arrow), clearly differentiated from the fully folded protein (F), is distinctly observable only in *CPk* thioredoxin. **b** Residue folding probabilities as a function of sequence index at 37°C following the colour code in panel **a**. **c** The predicted structure of F* with the partially structured residues 1-22 highlighted in orange. **d** Folding probability, coloured in the spectral scale from blue (0) to red (1), as a function of a plausible reaction coordinate, the number of structured blocks, for the three thioredoxin studied. The N-terminal region that folds the last is highlighted by white ovals while the 70-77 region is highlighted by an arrow. **e** Free energy landscapes (z-axis in kJ·mol^-1^) as function of n_N_ and n_C_, the number of structured blocks in the N- and C-terminal half of the protein, respectively, for the three thioredoxins studied here.

As mentioned above, thioredoxin folding has been known from many years to be a complex process that involves intermediate states and parallel channels to arrive to the native state^61,62^, reflecting a rugged folding landscape. We do not aim here at reproducing these kinetic complexities in detail, but mostly at identifying regions of the thioredoxin molecule that are likely to be unfolded in intermediate states of the folding landscape. The rationale behind this approach is that such unfolded or partially-structured regions may be involved in aggregation and other undesirable interactions^58^ and are, therefore, obvious targets for engineering efforts aimed at rescuing inefficient folding. This notion is consistent with early studies on the directed evolution of proteins for enhanced heterologous expression which demonstrated improvements in the stability and solubility of intermediates^69^.

We employ the block version of WSME model (the bWSME model), which considers 2-3 consecutive residues as blocks to reduce the protein phase space and thus rapidly calculate free energy profiles (see Methods). In the block description, the model still considers >490,000 microstates compared to the residue-level version that would involve considering the contribution to the partition function from >9.9 million microstates. We first reproduce the apparent experimental equilibrium stability differences to calibrate the model. The resulting one-dimensional folding free energy profiles as a function of the number of structured blocks, the reaction coordinate (RC), are quite similar for the three proteins, but with one major difference – *CPk* thioredoxin populates a partially structured intermediate (F*) on the folding side of the main barrier with a population of 27% (Fig. 4a). However, neither of LPBCA or *E. coli* thioredoxins populate F* significantly (<0.1%). It is important to note that this difference is not a consequence of the larger stability of *CPk* thioredoxin as *E. coli* thioredoxin does not populate F* despite exhibiting similar stability (black in Fig. 4a). Moreover, WSME model calculations reveal that the shape of the free-energy profiles is conserved under iso-stability conditions (of 25 kJ mol^-1^; Fig. S2). These observations indicate that the intermediate F* is intrinsic to the *CPk* thioredoxin conformational landscape and is independent of the overall thermodynamic stability.

To identify the regions of *CPk* thioredoxin that are partially structured, we computed residue folding probabilities that quantify the extent to which every residue is structured at a given thermodynamic condition. At 37°C, the N-terminal region of *CPk* thioredoxin (residues 1-22) is partially structured when compared to its ancestral counterpart (Fig. 4b). Differences in folding probabilities are also evident in two other regions: residues 70-77 that harbours a critical cis-proline^47,61^ and to a lesser extent in the residue stretch 83-84. We further calculated the probability of every region of the protein to be structured as a function of the reaction coordinate (Fig. 4d), and find that the N-terminal region of the protein folds the last, thus revealing the identity of F* (Fig 4c and white ovals in Fig. 4d). It can also be seen that the residue-stretch 70-77 exhibits equilibrium fluctuations during the folding for both the proteins (arrows in Fig. 4d). The two-dimensional free energy landscape (Fig. 4e), constructed by accumulating partial partition functions involving combinations of a given number of residues structured at N- and C-terminal halves of the protein, highlights that F* is the most populated state in *CPk* thioredoxin apart from multiple partially structured states that likely contribute to the slow folding of *CPk* thioredoxin (Figure 4e) compared to LPBCA thioredoxin.

There is a remarkable congruence between the computational predictions and the experimental folding kinetic data for *CPk* thioredoxin. Thus, a rate for reaching the native state much slower than expected from the typical shape of a chevron plot (compare continuous and discontinuous lines in the plot for *CPk* thioredoxin in Fig. 2a) is the pattern expected from the accumulation of an intermediate. Furthermore, the structure predicted for F* (Fig. 4c) is supported by results obtained with modern/ancestral chimeras described in the next section.

### Efficiency of heterologous expression in *E. coli* of modern/ancestral thioredoxin chimeras

On the basis of the folding-landscape computations described in the preceding section, we selected two regions as targets for the engineering of heterologous folding: the N-terminal 1-22 fragment, which includes a short *α*-helix and large stretches of non-regular structure, and the 70-77 loop which includes a cis-proline residue that has been known for many years to be critical for thioredoxin folding^47,61^. These two regions are predicted to fold comparatively late and to remain unfolded during most of the protein folding process (Fig. 4d). Plausibly, therefore, they may be involved in intermolecular processes that lead to insoluble protein. Since the heterologous expression of the ancestral LPBCA thioredoxin is efficient, we have prepared chimeras in which these regions are replaced by the corresponding ancestral sequences (see Fig. 5a). These replacements involve 17 mutational changes in the case of the 1-22 fragment and 4 mutational changes in the case of the 70-77 loop. We designate the chimeras as *CPk*-[1-22] thioredoxin and *CPk*-[70-77] thioredoxin. We have also studied the “double chimera” in which both regions are simultaneously replaced by the corresponding ancestral sequences: *CPk*-[1-22]-[70-77] thioredoxin. The three chimeras show substantially improved heterologous expression in *E. coli* with *CPk*-[1-22]-[70-77] thioredoxin approaching 100% of soluble protein (Fig. 5c and d).

**Figure 5.**
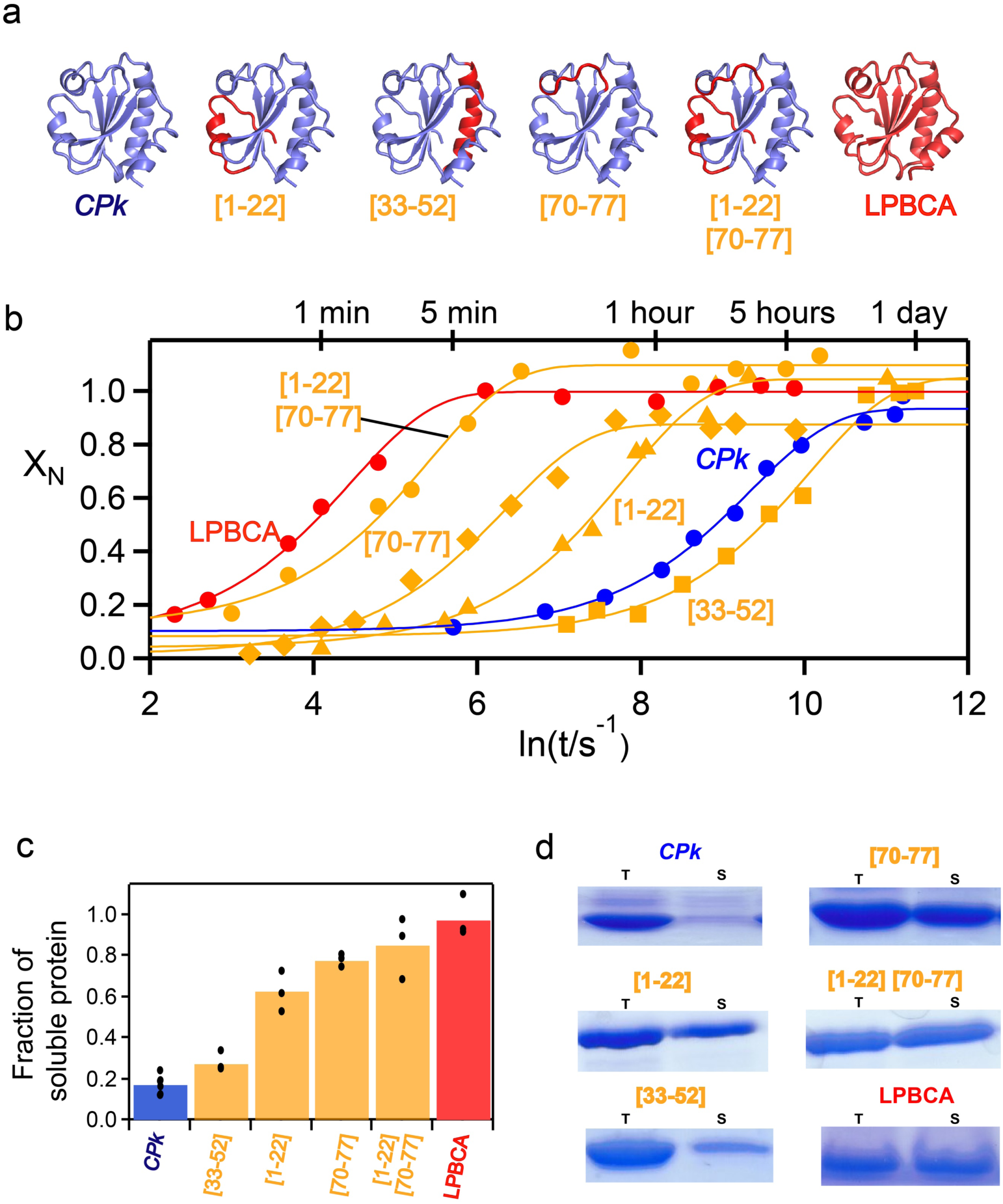
*In vitro* folding and *in vivo* expression of modern/ancestral chimeras. **a** Definition and structural description of the studied modern/ancestral chimeras. The thioredoxin backbone is coloured to indicate the origin of the sequence: modern *CPk* thioredoxin (blue) or ancestral LPBCA thioredoxin (red). **b** Profiles of fraction of native state *versus* time for the *in vitro* folding of *CPk* thioredoxin, LPBCA thioredoxin and the several modern/ancestral chimeras defined in **a**. The values of the fraction of native state are derived from double-jump unfolding assays (see Methods for details). The plot is labelled with characteristic time values to highlight the wide range of folding times for the proteins studied. The lines represent the best fits of a single exponential. **c** Fraction of soluble protein obtained upon expression in *E. coli* at 37°C of *CPk* thioredoxin, LPBCA thioredoxin and the four chimeras defined in **a**. Bars represent the average of several independent determinations and the individual values are also shown. **d** Representative examples of the experimental determination of the fraction of soluble protein for LPBCA thioredoxin, *CPk* thioredoxin and the modern/ancestral chimeras defined in **a**. Complete gels are shown Figures S10-S13.

In addition to the two regions referred to in the preceding paragraph, we have also selected for experimental analysis the 33-51 region which matches the longest *α*-helix in the thioredoxin molecule. This region is selected to provide an obvious control experiment, since it is predicted to fold early (Fig. 4d) and, according to our working hypothesis, we do not expect its replacement with the corresponding ancestral sequence to improve the efficiency of heterologous folding. The experimental results on *CPk*-[33-51] thioredoxin conform to this expectation (Fig. 5c and d).

### Efficiency of heterologous expression of single-mutant variants of the symbiont thioredoxin

As described in the preceding section, efficient heterologous expression of the symbiont thioredoxin is achieved through replacement of the 1-22 and 70-77 regions with the corresponding ancestral sequences in LPBCA thioredoxin. To explore the individual mutational contributions to the rescue, we have prepared 15 back-to-ancestor, single-mutant variants. These include the four back-to-ancestor mutations in the 70-77 region and 11 back-to-ancestor mutations in the 1-22 region. We have excluded from this analysis the initial 1-6 N-terminal segment, since it appears to be unstructured in the native structures. For all the 15 single-mutant variants we have determined the heterologous expression efficiency in *E. coli* (Fig.3a). The most remarkable result of these studies is that a single mutation at the 1-22 segment, S11N, rescues most of the inefficient heterologous folding. Combining this mutation with the best-rescuing mutation in the 70-77 loop leads to a double mutant variant of the symbiont thioredoxin, S11N/G74S, that approaches 100% soluble protein.

### Correlation between heterologous folding efficiency and the *in vitro* folding rate

The modern-ancestral chimeras, the single-mutant variants and the double S11N/G74S variant studied here are all active and show levels of *in vitro* redox activity similar to the modern *CPk* thioredoxin and *E. coli* thioredoxin, as well as the ancestral LPBCA thioredoxin (Fig. S1). They differ substantially, however, in terms of *in vitro* refolding rate. We have used double-jump unfolding assays to determine the rate constant for the last stage of *in vitro* refolding process, *i.e.,* the stage than leads to the native protein and defines the time scale of folding. These experiments reveal a very large (*∼*400-fold) range of folding rates (Figs. 2b, 5b and 6) with the time scale in which these proteins reach the native state *in vitro* varying between a few minutes and many hours. Remarkably, there is a good correlation between the efficiency of heterologous folding *in vivo* and the folding rate *in vitro* (Fig. 6a), supporting that efficient heterologous folding is achieved through the reduction of the time partially-unfolded states are significantly populated during the folding process.

**Figure 6.**
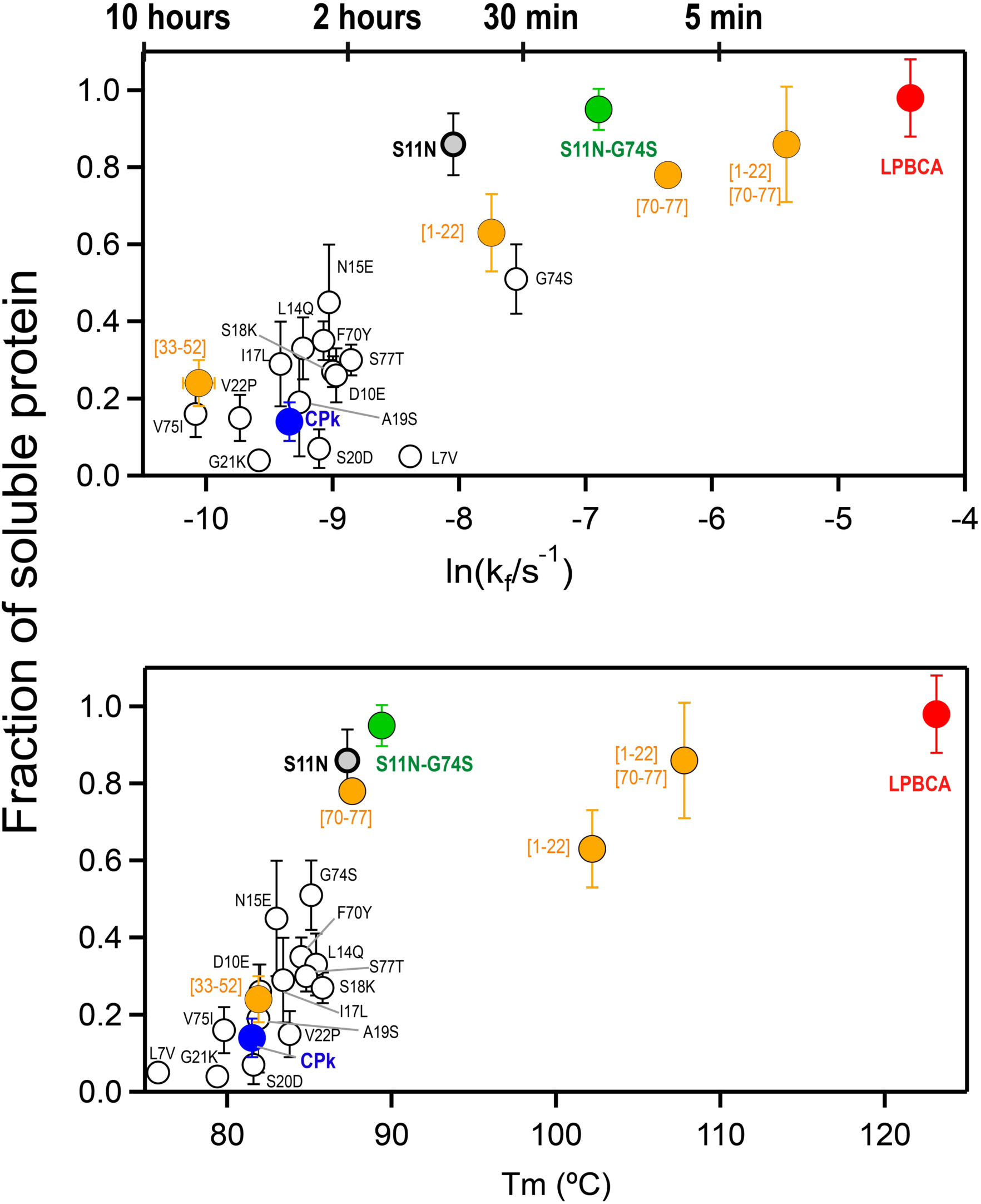
Correlations of the efficiency of *in vivo* heterologous expression with *in vitro* folding rate and protein stability. **a** Plot of fraction of soluble protein obtained in the expression in *E. coli versus* the logarithm of the *in vitro* folding rate constant including *CPk* thioredoxin, LPBCA thioredoxin, several variants of *CPk* thioredoxin and several modern/ancestral chimeras (Fig. 5). Typical values of the half-life time are indicated to highlight the wide range of folding time-scales. **b** Plot of fraction of soluble protein obtained in expression in *E. coli versus* denaturation temperature values derived from differential scanning calorimetry experiments. The proteins included here are the same as those included in the plot of panel **a**.

### Relation between heterologous folding efficiency and protein stability

We have used the denaturation temperature, as determined by differential scanning calorimetry, as a simple metric for the stability of the modern/ancestral chimeras, the single-mutant variants and the double S11N/G74S variant. For some selected variants, we have also performed urea denaturation studies. All the variants studied incorporate back-to-ancestor modifications. The ancestral sequence used as reference is that of LPBCA thioredoxin, a hyperstable protein with a very high denaturation temperature^48,60^. As anticipated, therefore, the back-to-ancestor modifications produce stability enhancements with respect to the *CPk* thioredoxin background in most cases. Furthermore, there appears to be a reasonable correlation between heterologous folding efficiency and stability, as described by the denaturation temperature values (Fig. 6b).

It is important to note, however, that global stability alone cannot explain the rescue of inefficient heterologous folding by back-to-ancestor modifications. This is more clearly seen when comparing the data of *CPk*-[1-22] thioredoxin with those for the variant of *CPk* thioredoxin with the single S11N mutation (Fig. 7). Both scanning calorimetry data and chemical denaturation profiles indicate that replacement of the 1-22 segment in *CPk* thioredoxin with the corresponding LPBCA ancestral sequence brings about a very large stabilization, while the single S11N mutation produces a much more moderate stability enhancement (Fig. 7a and b). Yet, heterologous folding of the single S11N variant is even more efficient than that of *CPk*-[1-22] thioredoxin (Fig. 7c). Obviously, much of the stabilization brought about by the replacement of the 1-22 segment in *CPk* thioredoxin with its ancestral counterpart has little effect on heterologous folding efficiency. Our results are, therefore, consistent with the notion that protein stabilization may improve heterologous expression^70^, but support that the rescuing effect of stabilization is linked to specific mutations in regions of the protein that are likely unfolded in aggregation-prone intermediate states.

**Figure 7.**
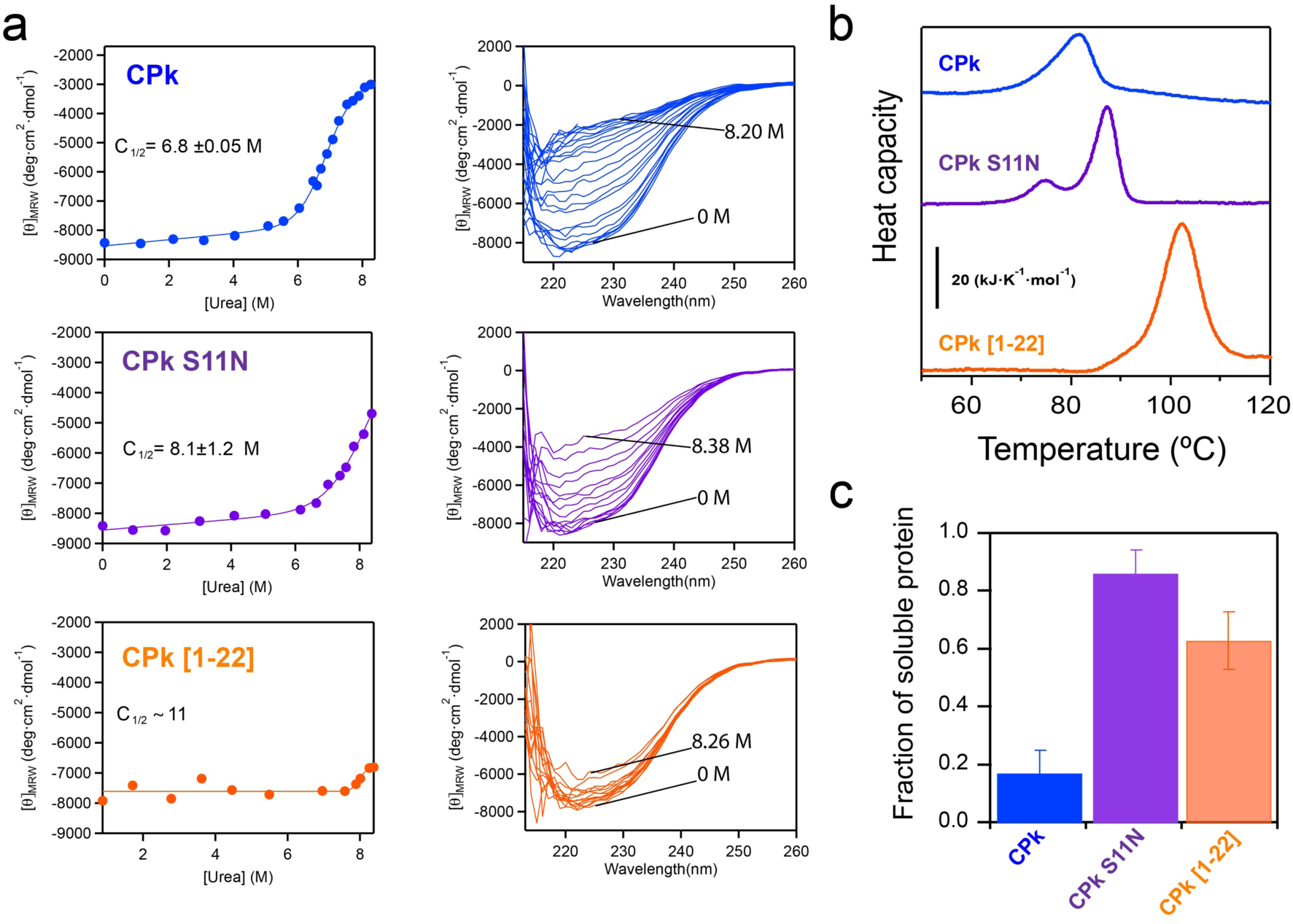
Relation between stabilization and rescue of inefficient heterologous expression. **a** and **b** Stability of the symbiont *CPk* thioredoxin, its S11N variant and the chimera in which the 1-22 segment has been replaced by the corresponding ancestral LPBCA sequence (*i.e., CPk*-[1-22] thioredoxin). In the experiments shown in **a** stability is probed by urea-induced denaturation followed by circular dichroism. Both, the original spectra at several urea concentrations and the profiles of ellipticity at 222 nm *versus* denaturant concentration are shown. The continuous lines represent the best fits of a two-state model (see Methods) and mid-point urea concentrations derived from the fits are shown. In **b** stability is studied by differential scanning calorimetry (see Methods and Fig. S15 for details). While the mutation S11N has a significant but moderate stabilizing effect, the chimera is highly stable as revealed by a high denaturation temperature and resistance to denaturation at 25°C by high urea concentrations. **c** Fraction of soluble protein obtained upon expression in *E. coli* at 37°C of *CPk* thioredoxin, its S11N variant and the *CPk*-[1-22] thioredoxin chimera. Despite its much-enhanced stability, the chimera is less efficient at rescuing heterologous expression than the S11N variant.

Finally, a somewhat intriguing result of our extensive experimental study on the stability of *CPk* thioredoxin variants deserves some attention. While most of the variants show the expected stability enhancement, this is not the case with *CPk*-[33-51] thioredoxin. This modern/ancestral chimera involves 10 back-to-ancestor amino acid replacements in the longest *α*-helix of the thioredoxin structure, which is expected to fold early according to our statistical-mechanical calculations (Fig. 4). *CPk*-[33-51] thioredoxin displays a stability similar to that of the *CPk* thioredoxin background as shown by both the denaturation temperature values and the chemical denaturation profiles (Fig. S3). One interesting possibility is that the ancestral stabilization is an adaptation to the need of efficient unassisted folding and high kinetic stability in an ancient environment. Consequently, it is not implemented in regions of the protein that are folded in aggregation-prone intermediates of the folding landscape and in the transition state that determines kinetic stability. Another possibility is that early folding of the long helix is crucial, mutations that impair its stability are not therefore accepted during evolution and, consequently, modern thioredoxins preserve the ancestral stability of the long helix. These interpretations are obviously speculative at this stage and will be explored in future work.

### Structural basis of the rescue of inefficient heterologous folding

Inefficient heterologous expression of *CPk* thioredoxin in *E. coli* is rescued to a substantial extent by a single S11N mutation. Ancestral sequence reconstruction is subject to uncertainties, but the posterior Bayesian probability for N at position 11 in the thioredoxin reconstruction corresponding to the last common ancestor of bacteria is close to unity^48^. As we have previously shown, sites with such high probability are very rarely, if ever, incorrectly predicted^71^. There is little doubt, therefore, that N is the ancestral residue at position 11 and that S11N is a back-to-ancestor mutation.

Position 11 is part of a type IV turn involving residues 8-11 (Fig. 8a). Turns are known to be crucial for protein folding in general, as they allow the polypeptide chain to fold onto itself and generate interactions that pertain to the native structure^72^. Presence of an asparagine residue at position 11 promotes the turn conformation because this residue can form stabilizing hydrogen bonds with the threonine at position 8 (Fig. 8b). On the other hand, a serine at position 11 is not predicted to form stabilizing hydrogen bonds with the threonine at position 8 (Fig. 8b) and consequently facilitates alternative conformations for the 8-11 segment in the high energy regions of the folding landscape. That is, the back to the ancestor mutation at position 11 should favour the local native conformation with respect local unfolded conformations, thus decreasing the time during which the corresponding partially unfolded states are significantly populated in the course of folding.

**Figure 8.**
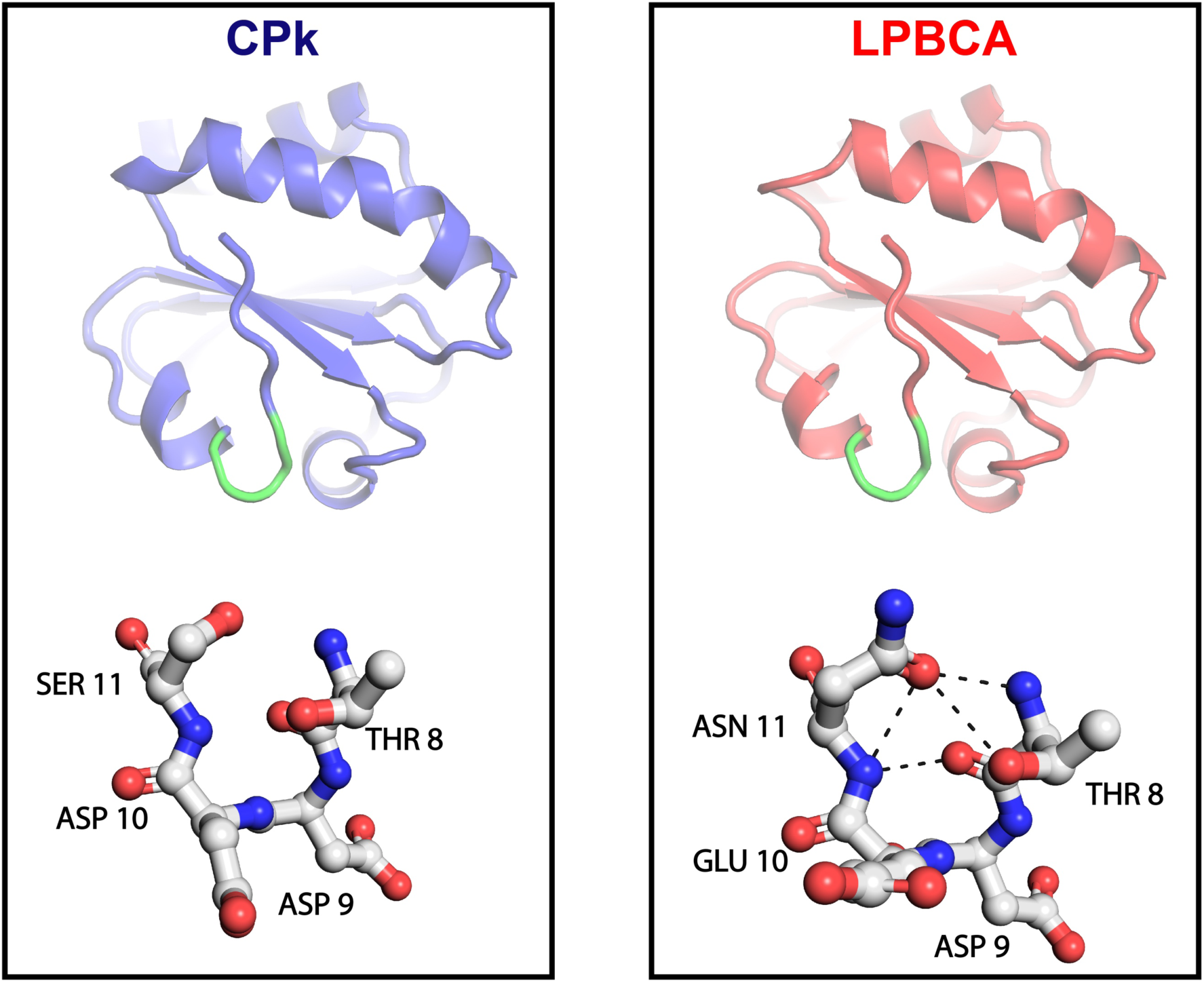
3D-structures of the modern *CPk* thioredoxin and the ancestral LPBCA thioredoxin highlighting a type IV turn including position 11. Highlights of the turn show the hydrogen bonds involving the residue in position 11 with other residues in the turn as predicted by WHAT IF^89^ with hydrogen-bond network optimization^90^. Hydrogen bonds are only predicted for the ancestral residue, which is therefore expected to stabilize the native turn conformation and disfavour non-native conformations.

A similar explanation can be deduced for the effect of the G74S mutation included in the S11N/G74S variant that approaches 100% soluble protein in heterologous expression. As we have previously noted^47^, effects on folding of the G/S exchange at position 74 in thioredoxins are very likely related the fact that glycine has no side chain and places little restriction in local backbone conformation. The flexible link generated by the presence of a glycine residue will allow many different conformations in the high energy region of the folding landscape. This is particularly relevant for the 70-79 loop, since it also includes the proline residue at position 76, which is in the rare cis-conformer in the native structure^61^. Presence of a glycine residue at position 74 thus enables many conformations for the 70-79 loop that are not consistent with the native cis conformation for Pro76.

## Concluding remarks

Our results support that the folding of proteins in heterologous hosts may be akin to some extent to unassisted folding. For the specific protein system studied here, this is supported by (i) the success of the approach used to rescue inefficient heterologous expression, which involved computational modelling of the unassisted folding landscape, (ii) the fact that the efficiency of heterologous expression correlates with the *in vitro* folding rate, (iii) the very moderate rescue of inefficient heterologous expression by chaperone over-expression.

Unassisted folding in heterologous hosts may conceivably result from the overexpression of the protein exceeding the capacity of the folding-assistance machinery. This is a reasonable scenario, in particular since co-evolution may have led to the adaptation of the protein to the assistance machinery of its original host. Consequently, the fact that resurrected ancestral proteins often show improved heterologous expression as compared with their modern counterparts^22–35^ plausibly reflects an ancient adaptation to unassisted folding. Efficient unassisted folding would no longer be a useful feature after the evolutionary emergence of cellular folding-assistance, thus allowing the evolutionary acceptance of mutations that impair the ancestral feature. Reversal of such mutations could then lead to a more efficient heterologous expression. Our results support that a few selected back-to-ancestor mutations can re-enact the folding efficiency of the resurrected ancestral proteins and point, therefore, to a minimal-perturbation, sequence-engineering approach to resolve inefficient heterologous expression. Some details of the practical application of the approach are noted below.

Prediction of back-to-ancestor mutations should be feasible for most protein systems, given the availability of large sequence databases and various software packages for the several steps of ancestral sequence reconstruction (for a recent account, see ref.^21^). Certainly, experimental screening for expression of all possible variants with single back-to-ancestor mutations may not be practical, in particular for large proteins. However, our results support that a limited screening of modern/ancestral chimeras may provide variants with enhanced expression. Furthermore, screening of individual mutations can be focused to protein regions that are expected to be unfolded in aggregation-prone intermediates populated during the folding process. Our results support that such regions can be predicted as the late-folding regions in folding landscape computations. Our version of the Wako-Saitô-Muñoz-Eaton statistical-mechanical model requires only a homology structure model as starting point and, most importantly, employs a block description to drastically reduce the number of microstates in the computation, thus allowing for a fast prediction even for large proteins. It is also worth noting that, for the protein system studied here, convincing molecular explanations can be put forward for the effect of the S11N and G74S mutations. This suggest the additional possibility of using rational design to determine the specific back-to-ancestor mutations that rescue inefficient heterologous folding.

Overall, our results open up the possibility of rescuing inefficient heterologous expression linked to low solubility by introducing a comparatively small number of back-to-ancestor mutations targeted to specific protein regions. In this way, the sequence would be minimally altered, and, likely, the properties of the encoded protein would be barely altered. This approach could be particularly useful in metagenomics studies that depend on sequence-based screening. Such studies identify enzymes having potential new activities of interest (pollutant or plastic degradation, for instance) based on the sequence similarity with known enzymes. In this scenario, achieving efficient expression with minimal sequence alteration will effectively contribute to the characterization of the new activity in the laboratory.

## Methods

### Expression and purification of thioredoxin variants

We followed procedures we have previously described in several publications^47–49,60^ with small modifications. Briefly, genes encoding *Candidatus Photodesmus katoptron* thioredoxin (*CPk*) and the *CPk* chimeras (*CPk* [1-22] and *CPk* [33-52]) were synthesized with a His-tag at the C-terminal and codon optimized for expression in *E. coli* cells. Mutations required for the single-mutant *CPk* variants (L7V, D10E, S11N, L14Q, N15E, I17L, S18K, A19S, S20D, G21K, V22P, F70Y, G74S, V75I, S77T), the double-mutant S11N/G74S variant and the chimera involving the loop [70-77] were introduced using the QuikChange Lighting Site-Directed Mutagenesis kit (Agilent Technologies) and the sequences were confirmed by DNA sequencing. Genes were cloned into pET24b(+) plasmid (GenScript Biotech) and transformed into *E. coli* BL21 (DE3) cells (Agilent). Protein expression was induced by 1 mM IPTG and cells were incubated overnight at 25°C in LB medium and. Cell pellets were sonicated and His-tagged proteins were purified using affinity chromatography (HisGraviTrap column from GE Healthcare).

*E. coli* thioredoxin, LPBCA thioredoxin and the G74S variant of the later used in the experiments reported in this work were prepared following procedures similar to those described above, except that His-tags were not used. Therefore, these proteins were purified^49^ by ion-exchange chromatography (Fractogel EMD DEAE column) followed by gel filtration chromatography on HiLoad Superdex 75 column. Our previous studies^47^ indicate that the presence of a His-tag has a very small effect on the folding kinetic features of thioredoxins. Also, the presence of a His-tag does not have a significant effect on the efficiency of heterologous expression for *CPk* thioredoxin. The purification procedure based on ion-exchange chromatography and gel filtration was also used to prepare the non-His-tagged *CPk* thioredoxin used for crystallization (see further below).

Folding-unfolding experiments reported in this work were performed with thioredoxin solutions in 50 mM Hepes, pH 7. These solutions were prepared either by dialysis against the buffer at 4°C or by passage through PD-10 desalting columns (GE Healthcare). Protein concentrations were measured spectrophotometrically using known values for the extinction coefficient. Guanidine and urea solutions in 50 mM HEPES, pH 7 were initially prepared by weight, but their concentrations were subsequently determined from refraction index measurements^73,74^ using an Atago R500 hand refractometer. Urea solutions were purified by passage of the stock solution through an AG501-X8(D) ion-exchange resin (Biorad) before use^74^.

### Double-jump unfolding assays

In principle, a protein in solution can populate a variety of conformational states, including, together with the native state, a diversity of unfolded and partially unfolded states. Hence, the value of a physical property for a protein solution may reflect contributions from different states and may potentially be difficult to interpret. Double-jump unfolding assays, on the other hand, provide an estimate of the amount of native state specifically^47,63,64^ and their interpretation is straightforward. They are based on the fact that the unfolding of the native state is much slower than the unfolding of intermediate (non-native and partially-unfolded) states. Therefore, the amount of native state in a protein solution can be estimated from the amplitude of the native-state unfolding kinetics observed upon transferring an aliquot to denaturing conditions, since intermediate states will unfold in a much shorter time-scale. Double-jump unfolding assays can be used to follow *in vitro* protein refolding kinetics (by performing the assays at several times during refolding), which provides an immediate assessment of the folding time-scale, *i.e.,* the time scale in which the folding polypeptide chain reaches the native state. This is particularly convenient when folding is a complex process involving multi-exponential kinetics and parallel kinetic channels, as is the case with thioredoxins^61,62^. We have recently described and discussed in some detail the use of double-jump unfolding assays to follow thioredoxin refolding kinetics^47^. Therefore, we provide below only the specific experimental details relevant to the experiments reported in this work.

In most cases, proteins were denatured in high urea concentration (within the range 7.5-9M) and, after fluorescence determinations had indicated that the unfolding process was essentially complete, the folding kinetics was initiated by dilution into native conditions (*i.e.,* into low denaturant concentration, typically 1M urea, although some experiments at other final urea concentrations were performed, as shown in panel a of Fig. 2). In most cases, the protein concentration in the folding kinetics experiments was on the order 0.1 mg/mL, although additional experiments at higher protein concentrations were carried out with *CPk* thioredoxin (see Fig. 2b). At different times after the start of a folding process, aliquots were extracted, transferred to denaturing conditions with typically a 1:15 dilution and the unfolding kinetics were determined by following the protein fluorescence at 350 nm as a function of time. The exact composition of the denaturing solution is immaterial for the result of the experiment, as long as the same composition is used in all the unfolding profiles corresponding to given folding experiment. Both, high urea concentrations (within the range 8-9.5 M) and high guanidine concentrations (within the range 3-5 M) were used. Of course, it is important that the denaturant concentration used does indeed unfolds the protein variant studied. In order to select denaturation concentrations that fulfill this criterion, we determined the unfolding branches of Chevron plots for all the thioredoxins studied here (Fig. S4). A few of the thioredoxins studied here are highly stable and they cannot be denatured by urea at 25°C, not even using the highest urea concentrations experimentally available. In these cases, the initial unfolding step was performed in concentrated guanidine (within the range 3-4.5 M) and the dilution into native conditions was designed to ensure a low guanidine concentration in the folding kinetics experiment (within the range 0.1-0.3 M). Figures S5-S8 show several representative examples of the experiments we have described. Folding kinetic profiles for all the proteins studied here are shown in Figs. 2b, 5b and S9.

In all cases, we carried out control experiments in which the native protein (at the same concentration used in the folding kinetic determinations) was transferred to the denaturing solution and the unfolding kinetics was followed by fluorescence at 350 nm. The amount of native protein at each time is then calculated as the ratio of the amplitude of the unfolding kinetics determined from the aliquot extracted at that time to the amplitude of the unfolding kinetics for the control. The resulting profiles of fraction of native state (X_N_) versus time conformed to a single exponential. Note, however, that the kinetic profiles shown in Figs. 2b, 5b and S9 use logarithm of time in the x-axis to highlight differences in folding time scale and that a single X_N_ *vs.* t exponential appears as sigmoidal in a plot of X_N_ *vs.* lnt.

Data of fraction of native protein *vs.* time were fitted with the following equation: X_N_=X*_∞_*+(X_0_-X*_∞_*)exp(-kt), where k is the first-order rate constant, and X_0_ and X*_∞_* are short-time and long-time limiting values of the fraction of native protein. Note that X_0_ and X*_∞_* do not necessarily equal zero and unity, respectively. Small differences with the control may certainly cause X*_∞_* to depart somewhat from unity. More importantly, thioredoxin folding is a complex process involving parallel kinetic channels and intermediate states^61,62^. A value of X_0_ significantly higher than zero might reflect a that a fraction of the molecules reaches the native state in a shorter time scale than that probed by our experiments (*i.e.,* a fast folding kinetic channel). Likewise, a value of X*_∞_* significantly smaller than unity that a fraction of the molecules reaches the native state in a longer time scale than that probed in our experiments (*i.e.,* a slow folding kinetic channel). In practice, however, the values of X_0_ and X*_∞_* determined from the fitting of the equation to our experimental folding profiles are reasonably close to 0 and 1 in essentially all cases. This implies that our experiments do identify the kinetic phase leading to the native state in the major folding channel. Therefore, we used as a metric of the folding time-scale the half-life time calculated from the rate constant value derived from the fittings (*i.e.,* 1/k).

### *In vivo* protein solubility measurements

*In vivo* solubility of overexpressed thioredoxins variants in *E. coli* BL21(DE3) strain was checked based on SDS-PAGE, following standard protocols. Briefly, at least 3 independent clones of each thioredoxin variant were grown up to an optical density of 0.6 and induced with 1mM IPTG for 3 hours at 37 °C. A 90 mL aliquot of the final culture was centrifuged at 4000 rpm, 10 min at 4 °C and the collected pellet was re-suspended in 6 mL of lysis buffer containing 20 mM Tris, pH 7.5, 50 mM NaCl and a protease inhibitor tablet (Roche cOmplete™). After sonication, two aliquots were taken. One aliquot was subjected to SDS-PAGE to estimate the total amount of protein. Other aliquot was centrifuged (15000 rpm for 10 min at 4 °C) and the supernatant was subjected to SDS-PAGE to provide the amount of soluble protein. The SDS-PAGE gels obtained in this work are shown in Figs. S10-S13.

Quantification of total and soluble thioredoxin fractions was carried out by SDS-PAGE on 15% Tris-glycine SDS-polyacrilamide gels and using ImageJ software (https://imagej.nih.gov/ij/) for image analysis of the thioredoxin bands stained by Coomassie dye. Illustrative densitometry profiles are given in Fig. S14. At least, three independent measurements were performed for each protein variant. The average value, the standard deviation and the individual values are given in Figs. 3 and 5.

In addition, we attempted to rescue the *in vivo* inefficient folding of *CPk* thioredoxin by co-overexpressing *E. coli* chaperones. Five plasmids designed to express the following “chaperone teams” were purchased from TAKARA Bio Inc: pG-KJE8 (expressing dnaK-dnaJ-grpE-groES-groEL), pGro7 (expressing groES-groEL), pKJE7 (expressing dnaK-dnaJ-grpE), pG-Tf2 (expressing groES-groEL-tig) and pTf16 (expressing the trigger factor). Chaperone plasmids were transformed into BL21(DE3) chemical competent cells containing the different thioredoxin plasmids.

### Activity measurements

Activity of thioredoxin proteins was measured using the insulin turbidimetric assay^75^ (Holmgren, 1979) as we have previously described^47^. Briefly, in this assay, disulfides reduction by dithiothreitol (DTT) catalysed by thioredoxin causes insulin aggregation, which is followed spectrophotometrically at 650 nm. The reaction mixture contains 0.1M phosphate buffer pH 6.5, 2 mM EDTA, 0.5 mg/mL of bovine pancreatic insulin and a final thioredoxin concentration of 1.5 μM. The reaction is initiated by addition of DTT to a 1 mM final concentration. Activity values for each variant reported were obtained from the maximum value of plots of dA_650nm_/dt *versus* time. A total of 3 independent measurements were carried out for each thioredoxin variant. The resulting average values and the corresponding standard deviations are reported in Fig. S1.

In addition, for some variants, thioredoxin activity was also assayed with thioredoxin reductase coupled to the reduction of DTNB^76^ as we have previously described^47^. Final conditions in the cuvette were: 0.05 M Tris-HCl, 2 mM EDTA pH 8, 0.05 mg/mL BSA, 0.5 mM DTNB, 0.25 mM NADPH and 0.15 μM *CPk* variants. Reaction was started by addition of thioredoxin reductase to a final concentration of 0.02 μM and monitored spectrophotometrically. A total of three independent measurements were performed for each thioredoxin variant. The resulting average values and the corresponding standard deviations are reported in Fig. S1.

### Urea-induced equilibrium denaturation monitored by CD and fluorescence measurements

The urea-induced equilibrium denaturation of *CPk* thioredoxin, its S11N single mutant and *CPk* [1-22] chimera was studied by using far-UV circular dichroism measurements at 25°C. Protein concentration was *∼*0.8 mg/mL in a 1mm cuvette. The urea dependence of ellipticity at 222 nm could be adequately fitted by a two-state model that assumes a linear dependence of the unfolding free energy with denaturant concentration within the transition region and linear pre- and post-transition baselines, as previously described^47^. Values of the midpoint urea concentration (C_1/2_) and the slope of the urea-dependence of the unfolding free energy (m) derived from these fits are given in Fig. 7.

### Denaturation Temperature measured by Differential Scanning Calorimetry

The denaturation temperatures of wild *CPk* thioredoxin and single- and double-mutant variants (L7V, D10E, S11N, L14Q, N15E, I17L, S18K, A19S, S20D, G21K, V22P, F70Y, G74S, V75I, S77T, S11N/G74S) and modern/ancestral chimeras (*CPk* [1-22], *CPk* [70-77], *CPk* [1-22]/[70-77], *CPk* [33-52]) were determined by differential scanning calorimetry as the temperature for maximum of the calorimetric transition. Experimental DSC thermograms are shown in Fig. S15. Note that, in a few cases, two transitions were reproducibly seen in the thermograms, perhaps reflecting a decreased unfolding cooperativity upon mutation. In these cases, the maximum of the major transition was used as a metric of thermal stability. The experiments were performed with a MicroCal Auto-PEAQ DSC calorimeter (Malvern), at pH 7 in 50 mM HEPES buffer. Typically, protein concentration was within the 0.4-0.6 mg/mL range and scan rate was 240 K/h. Standard protocols well established in our laboratory for thioredoxins were followed^48,77^. In all cases, protein solutions for the calorimetric experiments were prepared by exhaustive dialysis and the buffer from the last dialysis step was used in the reference cell of the calorimeter. Calorimetric cells were kept under excess pressure to prevent degassing during the scan. Several buffer-buffer baselines were recorded to ensure proper calorimeter equilibration prior to the protein run.

### Statistical-mechanical modeling of the folding landscape

The WSME model approach, its variants and parameterization are described elsewhere in detail^57^. Briefly, the model is Gō-like in its energetics requiring a starting structure and coarse-grains the polymer at the residue level. Thus, every residue is assumed to sample two sets of conformations – folded (represented as binary variable *1*) and unfolded (*0*) – contributing to a set of *2^N^* microstates for a *N*-residue protein. In the variant of the model used in the current work, we make two approximations to the original version^54,56^. First, we consider microstates restricted to only single-stretches of folded residues (single sequence approximation, SSA), two-stretches of folded residues (double sequence approximation, DSA) with no interaction across the island, and DSA with interaction across the structured islands if they interact in the native structure and even if the intervening residues are unfolded^78^. Second, we further reduce the accessible phase space by considering 2-3 consecutive residues to fold and unfold together (*i.e.* as a single block or unit). This model has been validated against the residue-level approximations and enables rapid predictions^53^. In addition, we include contributions from van der Waals interactions, all-to-all electrostatics^57^, simplified solvation terms and sequence and structure specific conformational entropy. At the time the simulations were performed, the 3D structure of *CPk* thioredoxin was not available. Therefore, homology modeling was employed to generate a model using the Robetta server^79^ and the *E. coli* thiorecoxin as a template. Still, the alpha carbon RMSD between the homology model and the experimental structure is 0.524 Å.

All simulations were performed at 37°C, pH 7.0 and 0.1 M ionic strength conditions with the following parameters: atomic-level interaction energy (ξ) of -70 J mol^-1^ for every heavy atom van der Waals interaction identified with a 6 Å spherical cut-off and excluding the nearest neighbours for LPBCA and *CPk* thioredoxin (−76 J mol^- 1^ for *E. coli* thioredoxin), change in heat capacity 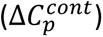 of -0.36 J mol^-1^ K^-1^ per native contact, entropic penalty for fixing non-proline, non-glycine and residues in well-determined secondary structures (Δ*S^conf^*) as -16.5 J mol^-1^ K^-1^ per residue, entropic penalty for glycine and coil residues as -22.56 J mol^-1^ K^-1^ per residue (accounting for the excess disorder in these regions^80^, and 0 J mol^-1^ K^-1^ per residue for proline residues given the limited backbone flexibility of prolines. Free energy profiles and surfaces as a function of the reaction coordinate, the number of structured blocks, are generated by accumulating partial partition functions corresponding to specific number of folded units. Residue folding probabilities are calculated by summing up the probabilities of states in which the residue of interest is structured.

### Crystallization and structural determination

Freshly purified *CPk* [70-77] thioredoxin was concentrated to 25 mg/ml prior setting the crystallization screening. Initial crystallization trials were carried out using the hanging-drop vapor diffusion method. Drops were prepared by mixing 1 µL of protein solution with the reservoir in a 1:1 ratio, and equilibrated against 500 1 µL of each precipitant cocktail of the HR-I & PEG/Ion™ crystallization screening (Hampton Research). Crystallization trials were kept at 293 k in an incubator. After one-week crystalline material was observed in conditions #33, #34, #43 and #11 of the HR-I kit and #13 of the PEG/Ion screening kit. Crystals were fished from the drop and transferred to the cryo-protectant solution prepared with the mother liquid supplemented with 15% (v/v) glycerol and subsequently flash-cooled in liquid nitrogen and stored until data collection.

Crystals were diffracted at ID23-1 beam-line of the European Synchrotron Radiation Facility (ESRF), Grenoble, France. The best diffracting crystals were obtained in condition #43 of the HR-I. The diffraction data were indexed and integrated using XDS^81^ and scaled with SCALA from the CCP4 suite^82^. Thioredoxin crystals belonged to the *I*2_1_3 space group with only two monomers in the asymmetric unit and therefore with a usually high water content, almost 74%, as determined from Matthewś coefficient, 4.71^83^. The molecular replacement solution was found using Molrep^84^ and the coordinates of the PDB ID. 2TRX, chain A, locating the two monomers in the asymmetric unit. Refinement was done with phenix.refine^85^ including manual building and water inspection with Coot^86^ and using the Titration-Libration-Screw (TLS)^87^ grouping since the initial steps. Model quality was checked using MolProbity^88^ implemented within the Phenix suite^85^. Refinement statistics and quality indicators of the final model are summarized in Table S1. Coordinates and structure factors have been deposited at the PDB with accession code 7OOL.

Hydrogen bond analysis was done using WHAT IF^89^ with hydrogen-bond network optimization^90^.

## Acknowledgements

This work was supported by Human Science Program Grant RGP0041/2017 (J.M.S.-R. and E.A.G.), National Science Foundation Award #2032315 (E.A.G.), National Institutes of Health Award #R01AR069137 (E.A.G.), Department of Defense MURI Award #W911NF-16-1-0372 (E.A.G.), Spanish Ministry of Science and Innovation/FEDER Funds Grants RTI-2018-097142-B-100 (J.M.S.-R.) and BIO2016-74875-P (J.A.G.) and the Science, Engineering and Research Board (SERB, India) Grant MTR/2019/000392 (A.N.N.). We are grateful to the European Synchrotron Radiation Facility (ESRF), Grenoble, France, for the provision of time and the staff at ID23-1 beamline for assistance during data collection.

## Author contributions

J.M.S.R. designed the research. G.G.-A. purified the modern/ancestral chimeras and the thioredoxin variants; she also performed and analysed the experiments aimed at determining their folding kinetics and biomolecular properties. V.A.R. performed experiments addressed at determining the efficiency of heterologous expression and provided essential input for the molecular interpretation of mutational effects on expression efficiency. E.A.G. provided essential input for the evolutionary interpretation of the data. J.A.G. determined the X-ray structure of the symbiont protein and provided essential input regarding its interpretation and implications. A.N.N. performed the computational simulations of the folding landscape for thioredoxins and provided essential input regarding their engineering implications. B.I.M. and J.M.S.-R. directed the project. J.M.S.-R. wrote the first draft of the manuscript to which V.A.R., J.A.G., A.N.N. and B.I.M. added crucial paragraphs and sections. All authors discussed the manuscript, suggested modifications and improvements, and contributed to the final version.

## SUPPLEMENTARY INFORMATION for

### Supplementary Tables

**Table S1.** Data collection and refinement statistics.

| Protein | Thioredoxin |
| --- | --- |
| PDB ID | 7OOL |
| Space group | I 21 3 |
| Unit cell |  |
| a, b, c (Å) | 140.08 140.08 140.08 |
| ASU | 2 |
| Resolution (Å) * | 44.3 - 2.85 (2.95 - 2.85) |
| Unique reflections | 10814 (1077) |
| Multiplicity | 3.5 (3.7) |
| Completeness (%) | 99.48 (99.91) |
| I/$\sigma_I$ * | 7.67 (1.03) |
| Wilson B-factor | 82.20 |
| R<sub>merge</sub> (%) * | 12.08 (93.51) |
| CC1/2 | 0.994 (0.454) |
| CC* | 0.998 (0.79) |
| **Refinement** |  |
| R<sub>work</sub>/R<sub>free</sub> (%) | 25.89 / 28.24 |
| No. atoms | 1749 |
| Protein | 1674 |
| Ligands | 63 |
| Solvent | 12 |
| B-factor (Å<sup>2</sup>) | 71.61 |
| RMS deviations |  |
| Bond lengths (Å) | 0.003 |
| Bond angles (°) | 0.69 |
| Ramachandran (%) |  |
| Favored | 99.04 |
| outliers (%) | 0.00 |
| Number of TLS groups | 2 |

\* Statistics for the highest-resolution shell are shown in parentheses.

**Table S2.** Percentage of sequence identity of chaperones from *Candidatus Photosmus kaptotron* with *E. coli* chaperones and with the closest hit from a BLAST search in the non-redundant protein sequences (nr) database using as query the sequence of *CPk* thioredoxin.

| Chaperone | Closest hit<br>from BLAST<br>search | *E. coli* |
| --- | --- | --- |
| Trigger factor | 51,3<br>*Vibrio maerlii* | 41,0 |
| groS | 91,7<br>*Vibrio sp E4404* | 79,3 |
| groEL | 91,5<br>*Vibrio tasmaniensis* | 84,2 |
| dnaK | 88,61<br>*Vibrio mexicanus* | 78,6 |
| dnaJ | 83,2<br>*Vibrio* | 68,9 |
| grpE | 79,2<br>*Vibrio parahaemolyticus* | 54,8 |

**Table S3.** Percentage of sequence identity of ribosomal proteins from *Candidatus Photosmus kaptotron* with *E. coli* ribosomal proteins and with the closest hit from a BLAST searches in the non-redundant protein sequences (nr) database using as query the sequences of the symbiont ribosomal proteins.

| **Ribosomal protein** | **Closest hit from BLAST search** | ***E. coli*** |
| --- | --- | --- |
| 30S ribosomal protein S1 (rpsA) | 85,5<br>*Vibrio alginolyticus* | 77,9 |
| 30S ribosomal protein S2 (rpsB) | 77,1<br>*Vibrio sp. 10N.261.55.A7* | 75,1 |
| 30S ribosomal protein S3 (rpsC) | 88,4<br>*Vibrio furnissii* NCTC 11218 | 82,9 |
| 30S ribosomal protein S4 (rpsD) | 82,5<br>*Vibrio europaeus* | 76,2 |
| 30S ribosomal protein S5 (rpsE) | 88,4<br>*Vibrio sp. 10N.261.55.A7* | 77,8 |
| 30S ribosomal protein S6 (rpsF) | 73,1<br>*Vibrio sp. 99-8-1* | 63,5 |
| 30S ribosomal protein S7 (rpsG) | 86,5<br>*Vibrio agarilyticus* | 78,8 |
| 30S ribosomal protein S8 (rpsH) | 77,7<br>*Vibrio cholerae* | 66,9 |
| 30S ribosomal protein S11 (rspK) | 94,2<br>*Vibrio parahaemolyticus*<br>EN9701121 | 84,5 |
| 30S ribosomal protein S9 (rpsI) | 85,8<br>*Vibrio* | 78,0 |
| 30S ribosomal protein S10 (rpsJ) | 100<br>*Vibrio cholerae* | 96,6 |
| 30S ribosomal protein S12 (rspL) | 86,6<br>*Vibrio parahaemolyticus* O1 | 90,3 |
| 30S ribosomal protein S13 (rspM) | 84,8<br>*Vibrio sp. HA2012* | 78,0 |
| 30S ribosomal protein S14 (rspN) | 82,2<br>*Candidatus Photodesmus blepharus* | 71,3 |
| 30S ribosomal protein S15 (rpsO) | 76,4<br>*Vibrio alginolyticus* | 65,2 |
| 30S ribosomal protein S16 (rpsP) | 67,5<br>*Vibrio* | 61,2 |
| 30S ribosomal protein S17 (rpsQ) | 79,7<br>*Vibrio sp. Hep-1b-8* | 69,2 |
| 30S ribosomal protein S18 (rspR) | 89,2<br>*Shewanella frigidimarina* | 86,5 |
| 30S ribosomal protein S19 (rspS) | 92,4<br>*Vibrio sp. ECSMB14106* | 84,8 |
| 30S ribosomal protein S20 (rspT) | 80,2<br>*Vibrio sp. V28_P6S34P95* | 64,4 |
| 30S ribosomal protein S21 (rspU) | 94,4<br>*Aeromonadales bacterium* | 85,9 |
| 50S Ribosomal protein L1 (rplA) | 66,4<br>*Alteromonas sp. KUL42* | 62,6 |
| 50S Ribosomal protein L2 (rplB) | 84,7<br>*Vibrio sp. qd031* | 75,9 |
| 50S Ribosomal protein L3 (rplC) | 77,9 | 69,7 |
|  | *Vibrio sp. S17_S38* |  |
| 50S Ribosomal protein L4 (rplD) | 77,9<br>*Vibrio inusitatus* NBRC 102082 | 67,7 |
| 50S Ribosomal protein L5 (rplE) | 83,8<br>*Vibrio alginivorius* | 76,0 |
| 50S Ribosomal protein L6 (rplF) | 66,9<br>*Vibrio hepatarius* | 59,6 |
| 50S Ribosomal protein L8 (rplI) | 67,1<br>*Vibrio natriegens* | 58,8 |
| 50S Ribosomal protein L9 (rplJ) | 72,5<br>*Vibrio parahaemolyticus* | 60,7 |
| 50S Ribosomal protein L10 (rplK) | 82,6<br>*Salinivibrio sp. KP-1* | 73,9 |
| 50S Ribosomal protein L7/ L12 (rplL) | 71,1<br>*Vibrio aestuarianus subsp. francensis* | 64,5 |
| 50S Ribosomal protein L13 (rplM) | 76,1<br>*Vibrio marinisediminis* | 62,0 |
| 50S Ribosomal protein L14 (rplN) | 89,5<br>*Vibrio parahaemolyticus* | 80,6 |
| 50S Ribosomal protein L15 (rplO) | 76,2<br>*Vibrio sp. 10N.286.49.C2* | 63,6 |
| 50S Ribosomal protein L16 (rplP) | 89,0<br>*Vibrio parahaemolyticus* | 79,4 |
| 50S Ribosomal protein L17 (rplQ) | 76,7<br>*Salmonella enterica subsp. enterica serovar Newport* | 75,4 |
| 50S Ribosomal protein L18 (rplR) | 77,1<br>*Vibrio galathea* | 64,4 |
| 50S Ribosomal protein L19 (rplS) | 87,1<br>*Vibrio sp. V37_P2S8PM304* | 74,1 |
| 50S Ribosomal protein L20 (rplT) | 85,1<br>*Aliivibrio salmonicida LFI1238* | 81,2 |
| 50S Ribosomal protein L21 (rplU) | 77,7<br>*Marinifaba aquimaris* | 65,7 |
| 50S Ribosomal protein L22 (rplV) | 85,5<br>*Vibrio tubiashii ATCC 19109* | 69,1 |
| 50S Ribosomal protein L23 (rplW) | 73,4<br>*Salmonella enterica subsp. enterica serovar Kentucky* | 60,6 |
| 50S Ribosomal protein L24 (rplX) | 76,2<br>*Vibrio cincinnatiensis DSM 19608* | 69,9 |
| 50S Ribosomal protein L25 (rplY) | 66,3<br>*Vibrio sp. S11_S32* | 45,3 |
| 50S Ribosomal protein L27 (rpmA) | 88,1<br>*Shewanella sp. Scap07* | 84,5 |
| 50S Ribosomal protein L28 (rpmB) | 87,5<br>*Vibrio parahaemolyticus* | 76,3 |
| 50S Ribosomal protein L29 (rpmC) | 72,6<br>*Rheinheimera sp. SA_1* | 61,9 |
| 50S Ribosomal protein L30 (rpmD) | 82,8<br>*Vibrio hepatarius* | 69,6 |
| 50S Ribosomal protein L32 (rpmF) | 83,9<br>*Vibrio* | 68,4 |
| 50S Ribosomal protein L33 (rpmG) | 81,8<br>*Candidatus Enterovibrio luxaltus* | 69,1 |
| 50S Ribosomal protein L34 (rpmH) | 87,5<br>*Vibrio parahaemolyticus 49* | 69,8 |
| 50S Ribosomal<br>protein L35 (rpml) | 89,7<br>*Vibrio sp. Y2-5* | 54,5 |
| 50S Ribosomal<br>protein L36 (rpmJ) | 92,0<br>*Glaciecola sp. 33A* | 73,7 |

**Figure S1.**
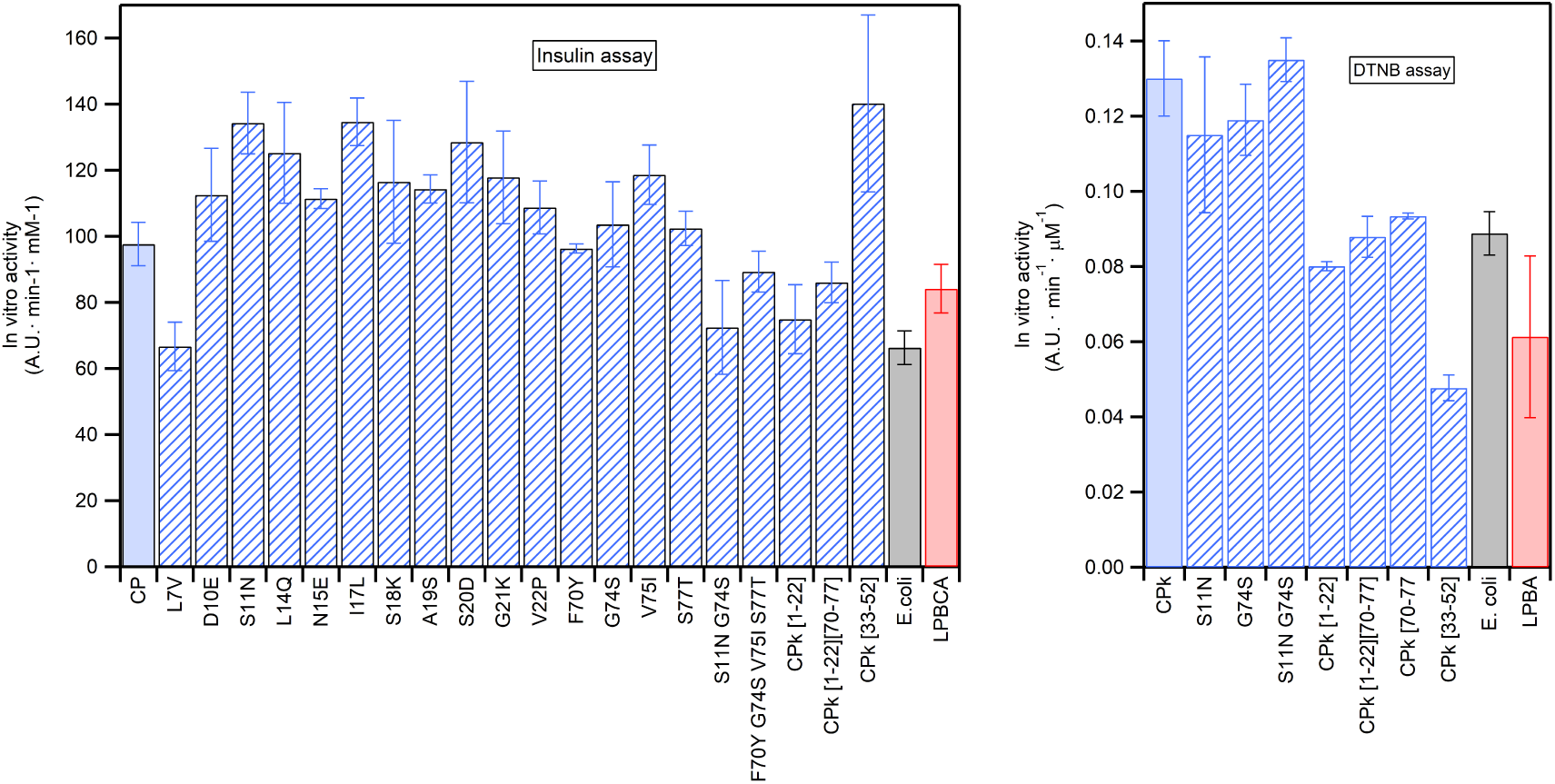
*In vitro* activities of thioredoxins studied in this work at 37 ªC. We used the insulin turbidimetric assay (left) and the assay with thioredoxin reductase coupled to the reduction of DTNB (right). The values given are the average of at least 3 independent measurements and the error bars represent the corresponding standard deviations.

**Figure S2.**
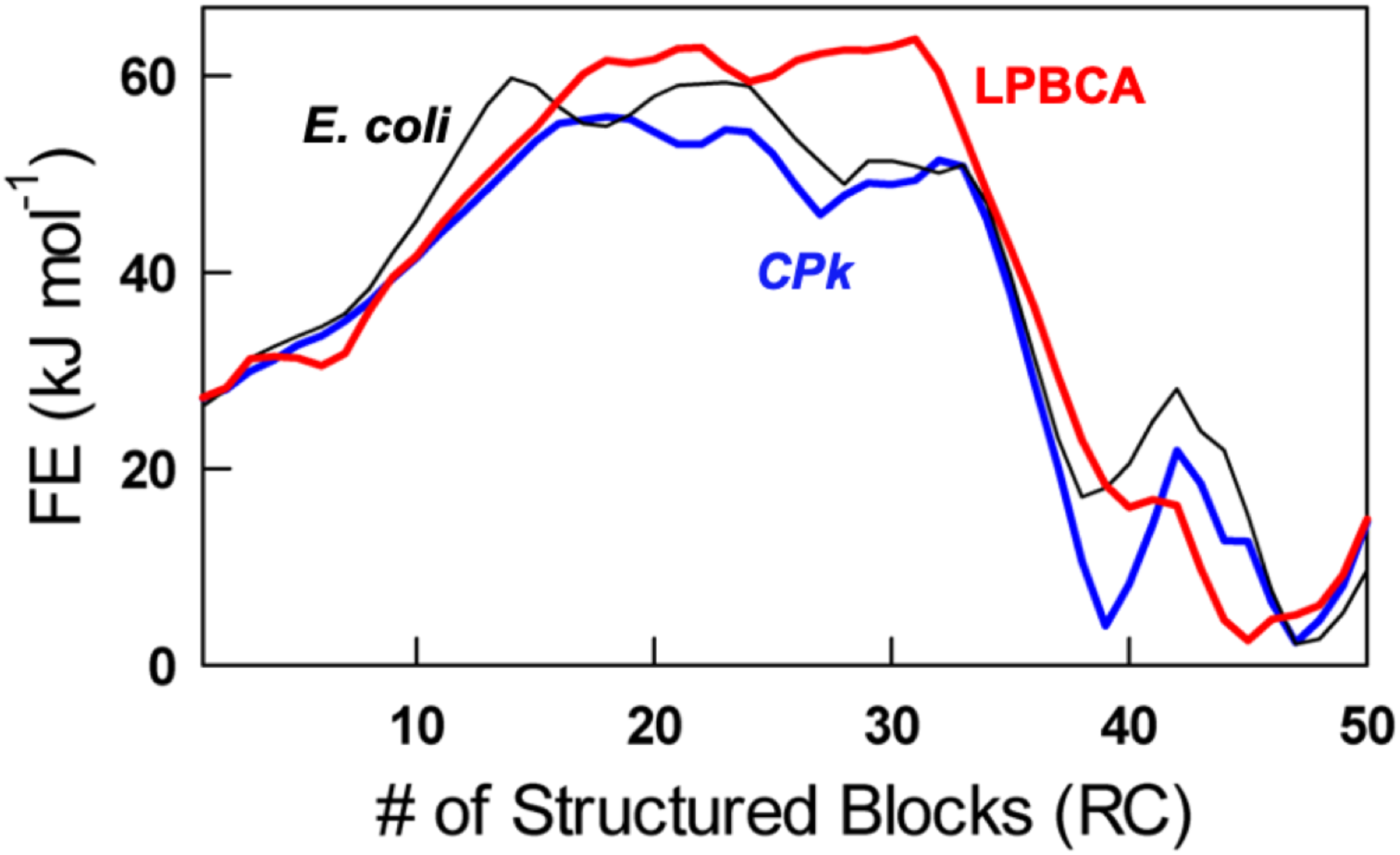
Profiles of free energy versus number of structured blocks at 37 °C for the modern thiorredoxins from *E. coli* and *Candidatus Photodesmus kaptotron* (*CPk*), and the ancestral LPBCA thioredoxin. These profiles have been calculated using the WSME model under iso-stability conditions (25 kJ·mol^-1^).

**Figure S3.**
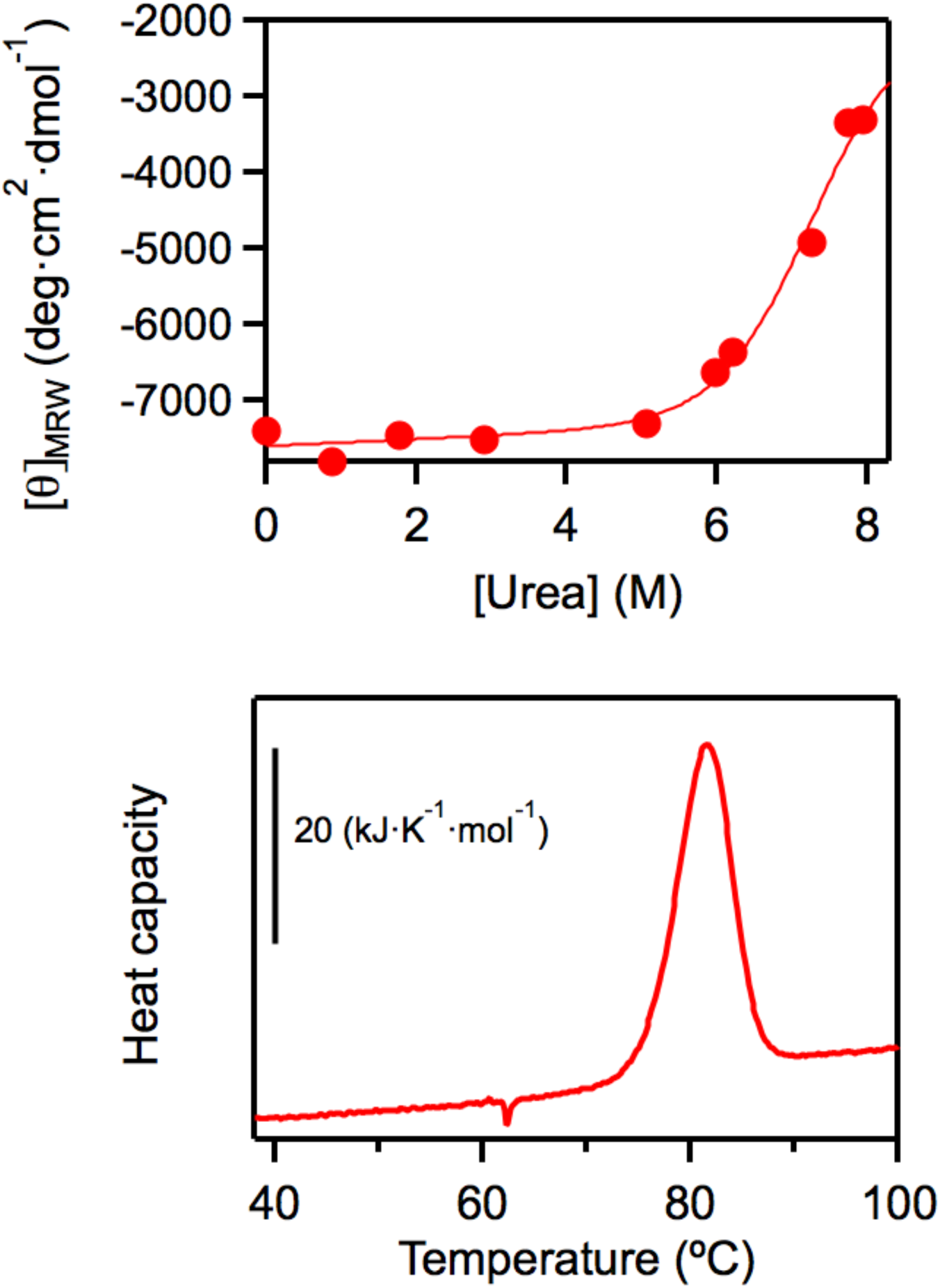
Stability of the *CPk*-[33-52] thioredoxin, a modern/ancestral chimera, as probed by urea-denaturation followed by circular dichroism and by differential scanning calorimetry. These experiments show a stability for *CPk*-[33-52] thioredoxin similar to that of *CPk* thioredoxin.

**Figure S4.**
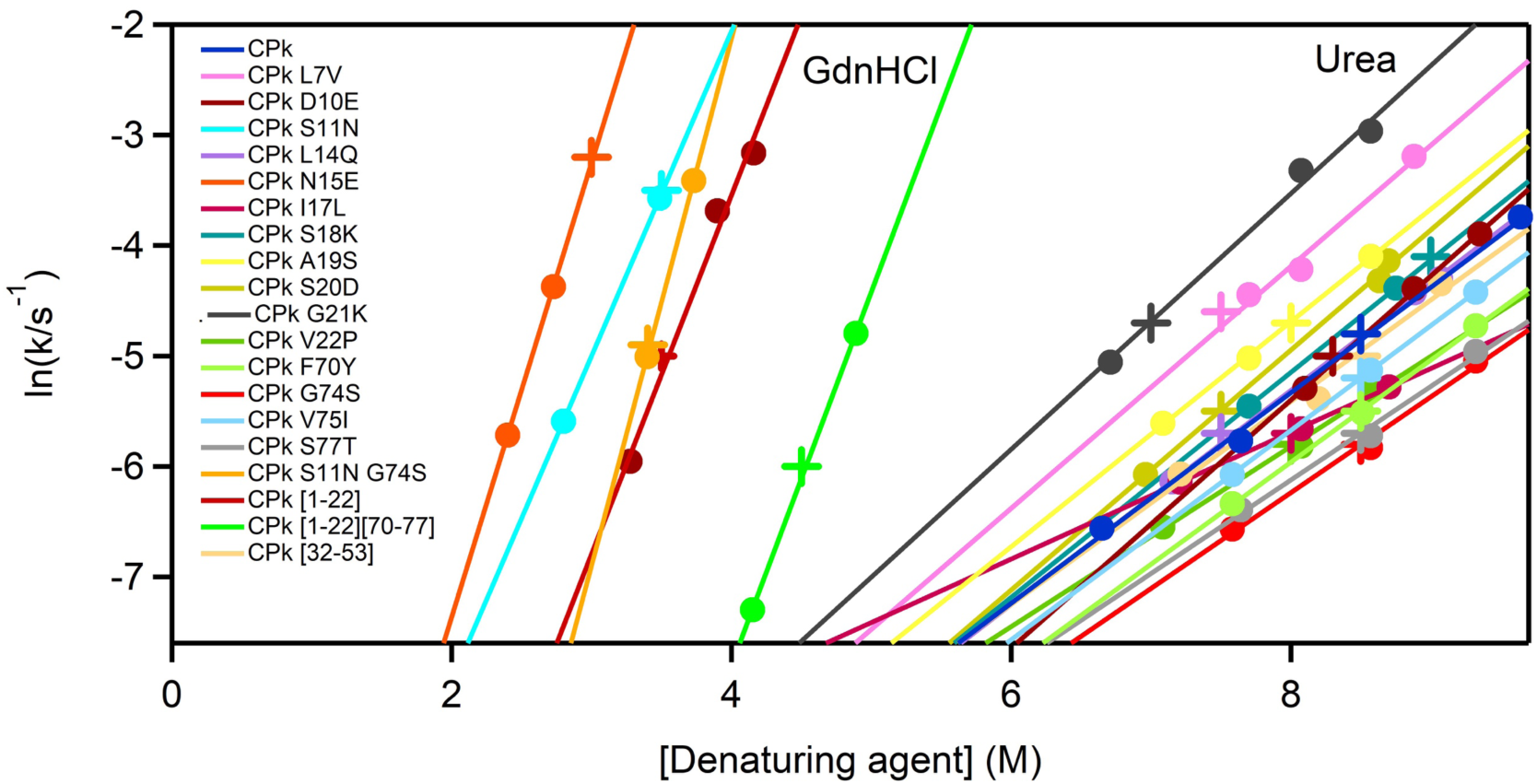
Unfolding branches of Chevron plots at pH 7 and 25 °C for thioredoxins studied in this work. The unfolding rate constants were determined from the fluorescence kinetic profiles obtained after transferring the native protein to concentrated denaturant solution. The purpose of this experiments was to determine denaturant concentrations suitable for the double-jump unfolding assays. Therefore, only a limited number of experiments were performed for each protein. The plus signs identify the denaturant concentration used to unfold the protein in the first step of the double-jump unfolding assays.

**Figure S5.**
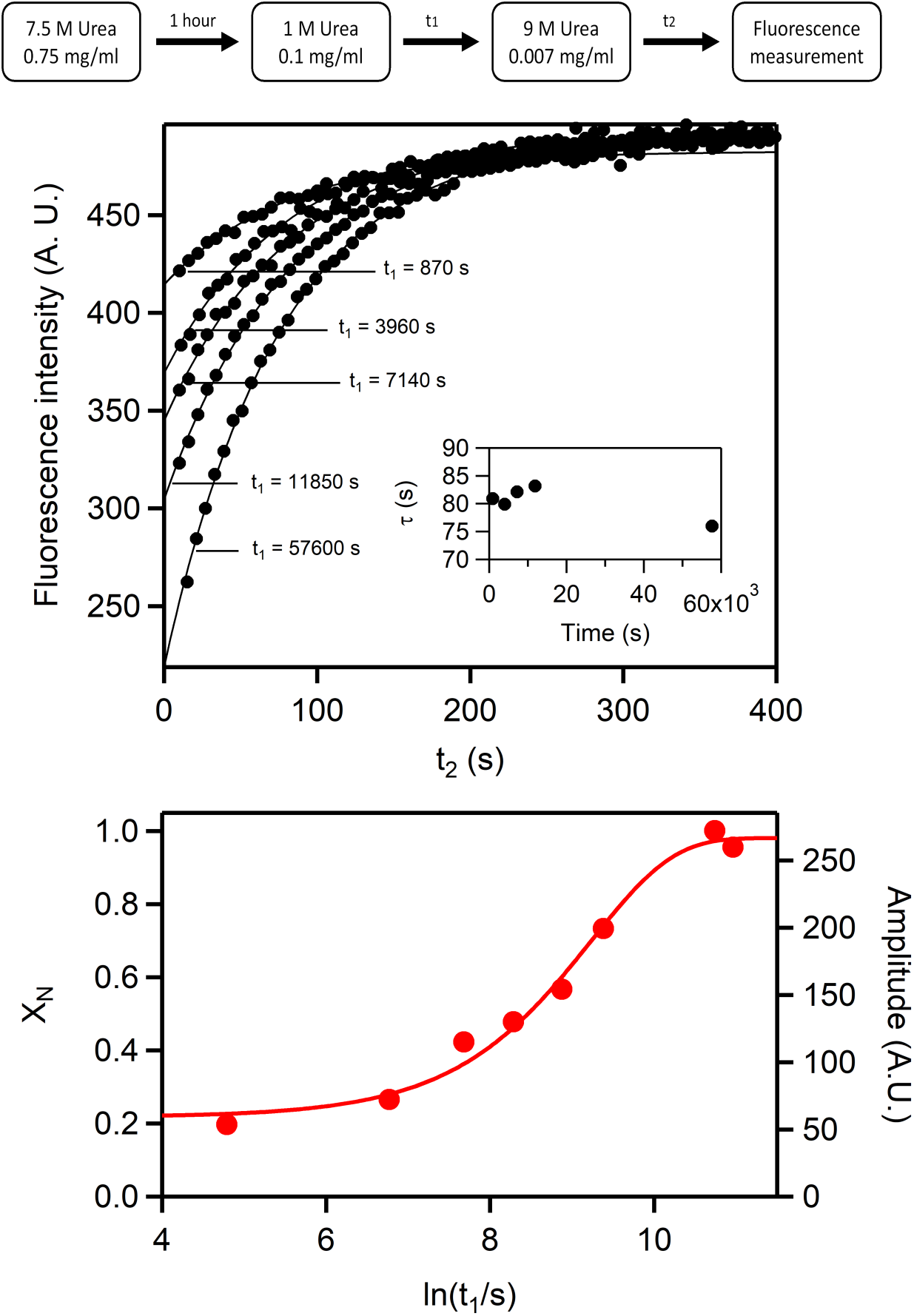
Representative example of a folding kinetics followed by double-jump unfolding assays. The data correspond to a variant of *CPk* thioredoxin with the single L14Q mutation. **a** Folding was initiated by transferring protein unfolded in 7.5 M urea to 1M urea. At different t_1_ times aliquots were transferred to denaturing conditions (9M urea) and the unfolding kinetics was followed by measuring fluorescence as a function of time (t_2_). **b** Plots of fluoresce intensity versus t_2_ time for unfolding, corresponding to different values of the t_1_ time. The continuous lines represent the best fits of a single exponential to the experimental data. These kinetic profiles are described essentially by the same unfolding rate constant (corresponding to 9M urea), as shown in the Inset. The amplitudes, however, do change, reflecting the folding kinetics in 1M urea and they lead to estimates of the fraction of native protein when normalized by the amplitude obtained in a control experiment. **c** Profile of fraction of native protein *versus* time for the folding in 1M urea. The line represents the best fit of a single exponential. Note that an exponential has a sigmoidal shape when the logarithm of the independent variable is used in the x-axis.

**Figure S6.**
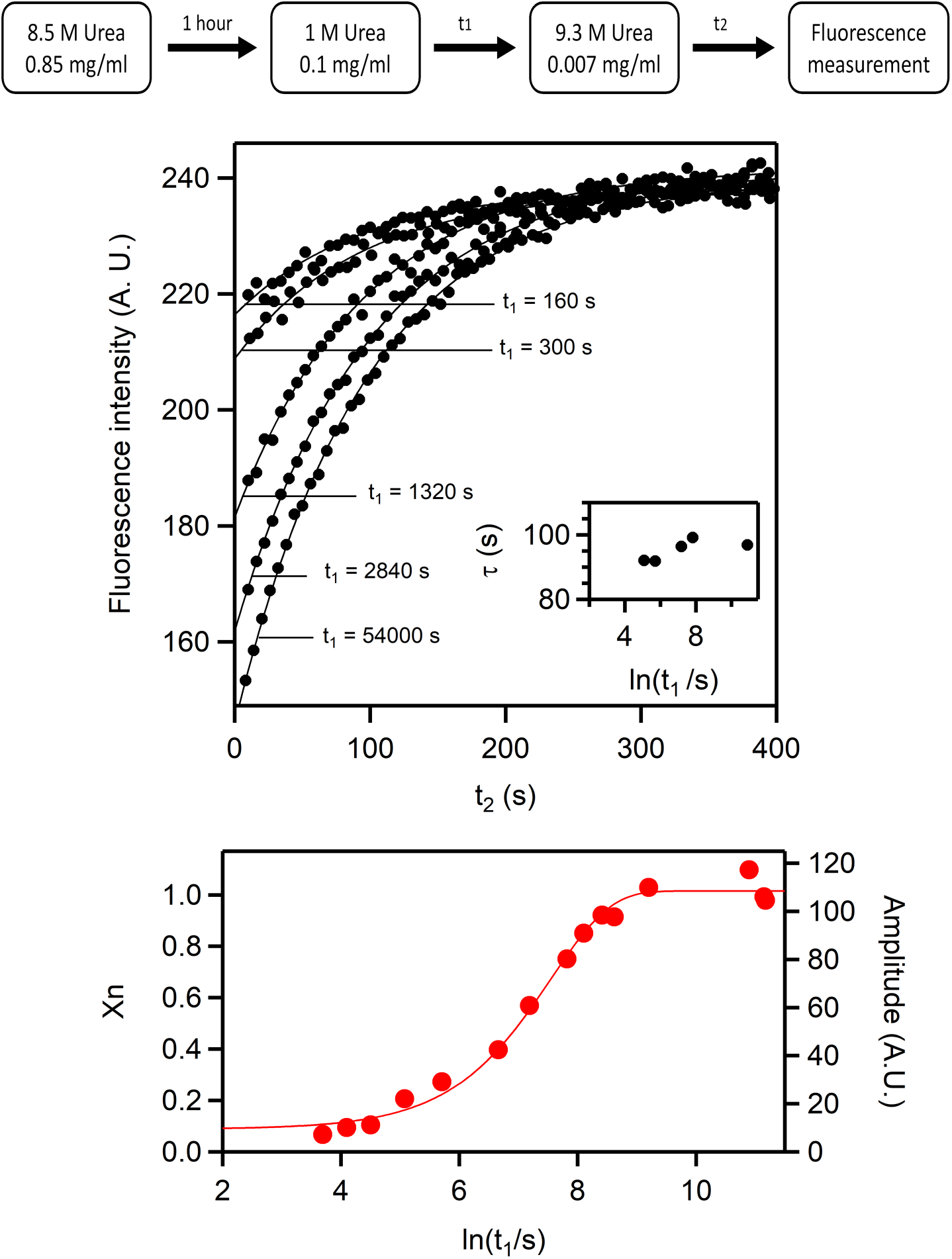
Representative example of a folding kinetics followed by double-jump unfolding assays. The data correspond to a variant of *CPk* thioredoxin with the single G74S mutation. **a** Folding was initiated by transferring protein unfolded in 8.5 M urea to 1M urea. At different t_1_ times aliquots were transferred to denaturing conditions (9.3 M urea) and the unfolding kinetics was followed by measuring fluorescence as a function of time (t_2_). **b** Plots of fluoresce intensity versus t_2_ time for unfolding, corresponding to different values of the t_1_ time. The continuous lines represent the best fits of a single exponential to the experimental data. These kinetic profiles are described essentially by the same unfolding rate constant (corresponding to 9.3 M urea), as shown in the Inset. The amplitudes, however, do change, reflecting the folding kinetics in 1M urea and they lead to estimates of the fraction of native protein when normalized by the amplitude obtained in a control experiment. **c** Profile of fraction of native protein *versus* time for the folding in 1M urea. The line represents the best fit of a single exponential. Note that an exponential has a sigmoidal shape when the logarithm of the independent variable is used in the x-axis.

**Figure S7.**
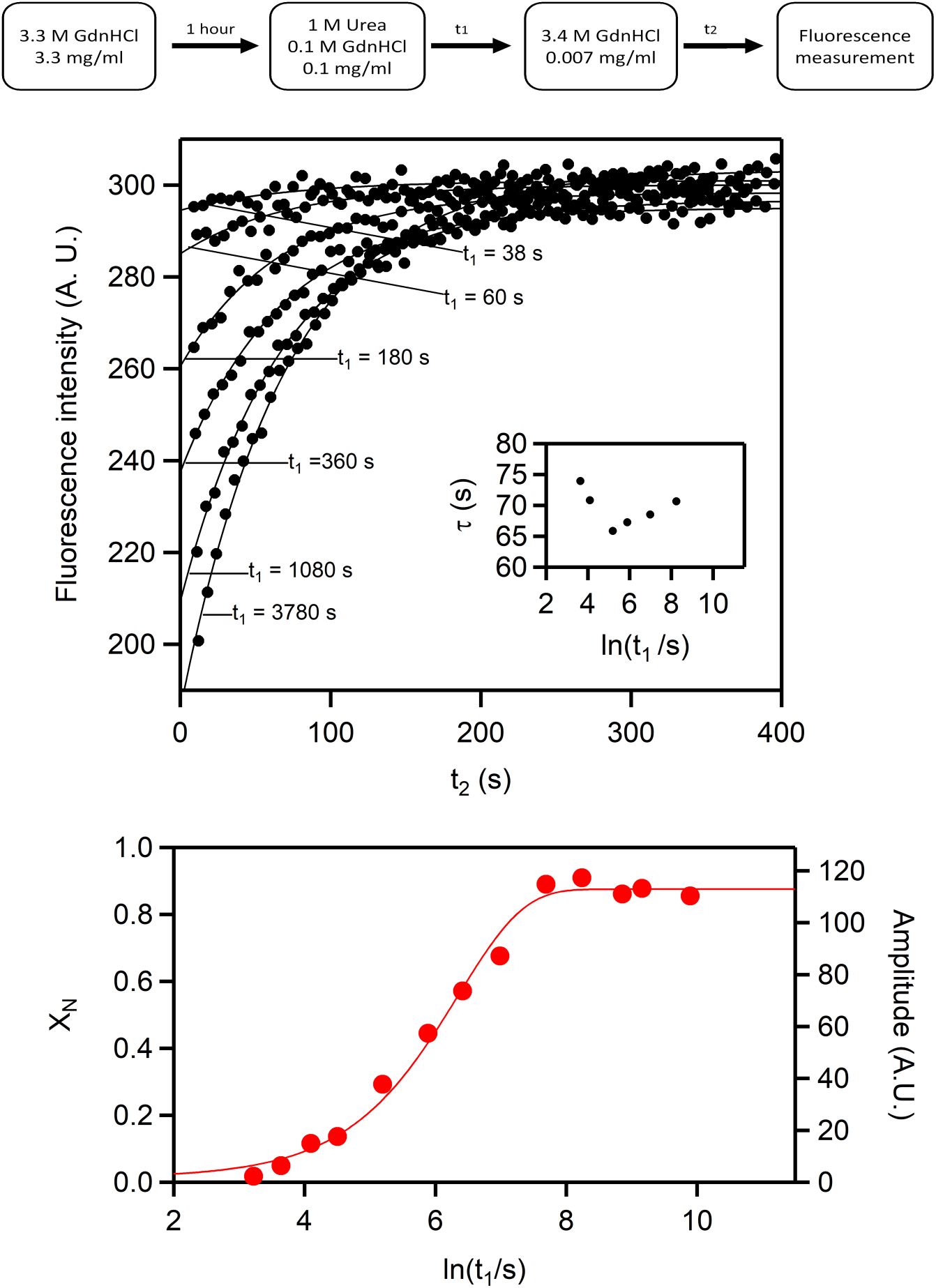
Representative example of a folding kinetics followed by double-jump unfolding assays. The data correspond to a the modern/ancestral chimera *CPk-*[70-77] thioredoxin with the single L14Q mutation. **a** Folding was initiated by transferring protein unfolded in 3.3 M guanidine to 1M urea. At different t_1_ times aliquots were transferred to denaturing conditions (3.4 M guanidine) and the unfolding kinetics was followed by measuring fluorescence as a function of time (t_2_). **b** Plots of fluoresce intensity versus t_2_ time for unfolding, corresponding to different values of the t_1_ time. The continuous lines represent the best fits of a single exponential to the experimental data. These kinetic profiles are described essentially by the same unfolding rate constant (corresponding to 3.4 M urea), as shown in the Inset. The amplitudes, however, do change, reflecting the folding kinetics in 1M urea and they lead to estimates of the fraction of native protein when normalized by the amplitude obtained in a control experiment. **c** Profile of fraction of native protein *versus* time for the folding in 1M urea. The line represents the best fit of a single exponential. Note that an exponential has a sigmoidal shape when the logarithm of the independent variable is used in the x-axis.

**Figure S8.**
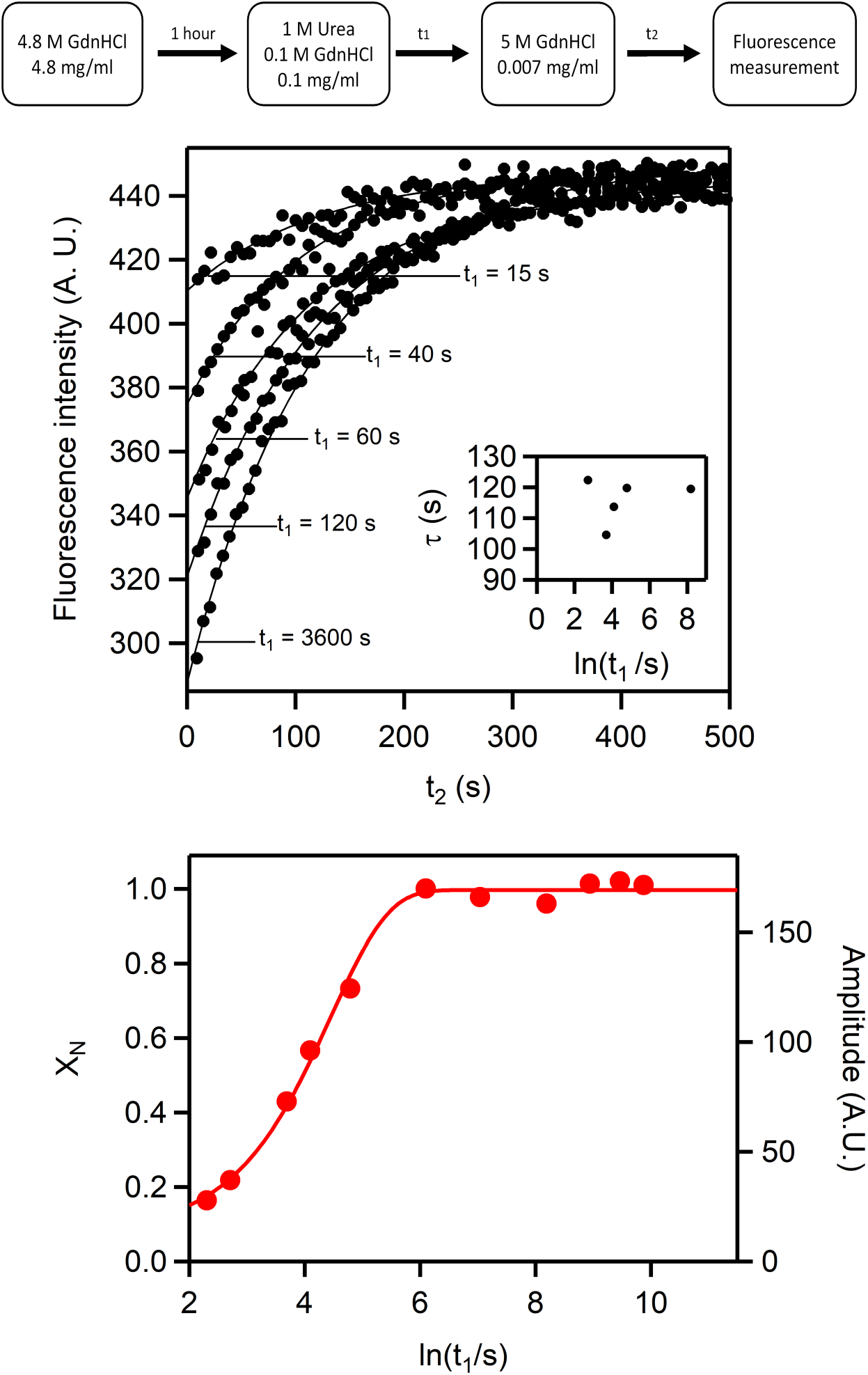
Representative example of a folding kinetics followed by double-jump unfolding assays. The data correspond to the ancestral LPBCA thioredoxin.**a** Folding was initiated by transferring protein unfolded in 4.8 M guanidine to 1M urea. At different t_1_ times aliquots were transferred to denaturing conditions (5M guanidine) and the unfolding kinetics was followed by measuring fluorescence as a function of time (t_2_). **b** Plots of fluoresce intensity versus t_2_ time for unfolding, corresponding to different values of the t_1_ time. The continuous lines represent the best fits of a single exponential to the experimental data. These kinetic profiles are described essentially by the same unfolding rate constant (corresponding to 5M guanidine), as shown in the Inset. The amplitudes, however, do change, reflecting the folding kinetics in 1M urea and they lead to estimates of the fraction of native protein when normalized by the amplitude obtained in a control experiment. **c** Profile of fraction of native protein *versus* time for the folding in 1M urea. The line represents the best fit of a single exponential. Note that an exponential has a sigmoidal shape when the logarithm of the independent variable is used in the x-axis.

**Fgure S9.**
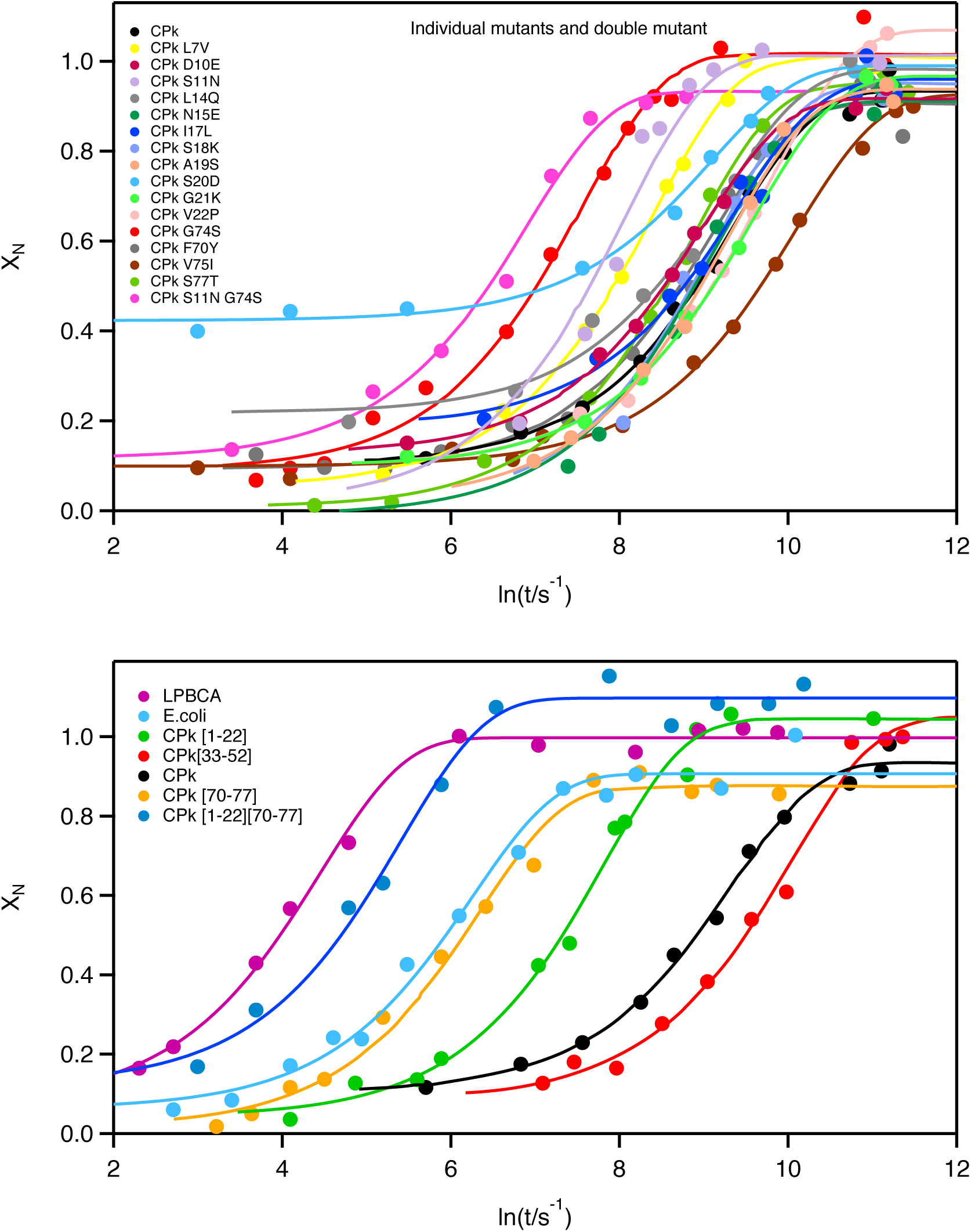
Folding kinetics at pH 7 and 1M urea, as followed by double-jump unfolding assays for all the thioredoxins studied in this work. The continuous lines represent the best fits of a single exponential to the experimental profiles of fraction of native protein *versus* time. Note that an exponential has a sigmoidal shape when the logarithm of the independent variable is used in the x-axis. With the exception of the S20D variant of *CPk* thioredoxin, in all cases the short-time and long-time limits are reasonably close to zero and unity, respectively, indicating that the analysis has identified the kinetic phase that leads to the native protein in the major folding channel.

**Figure S10.**
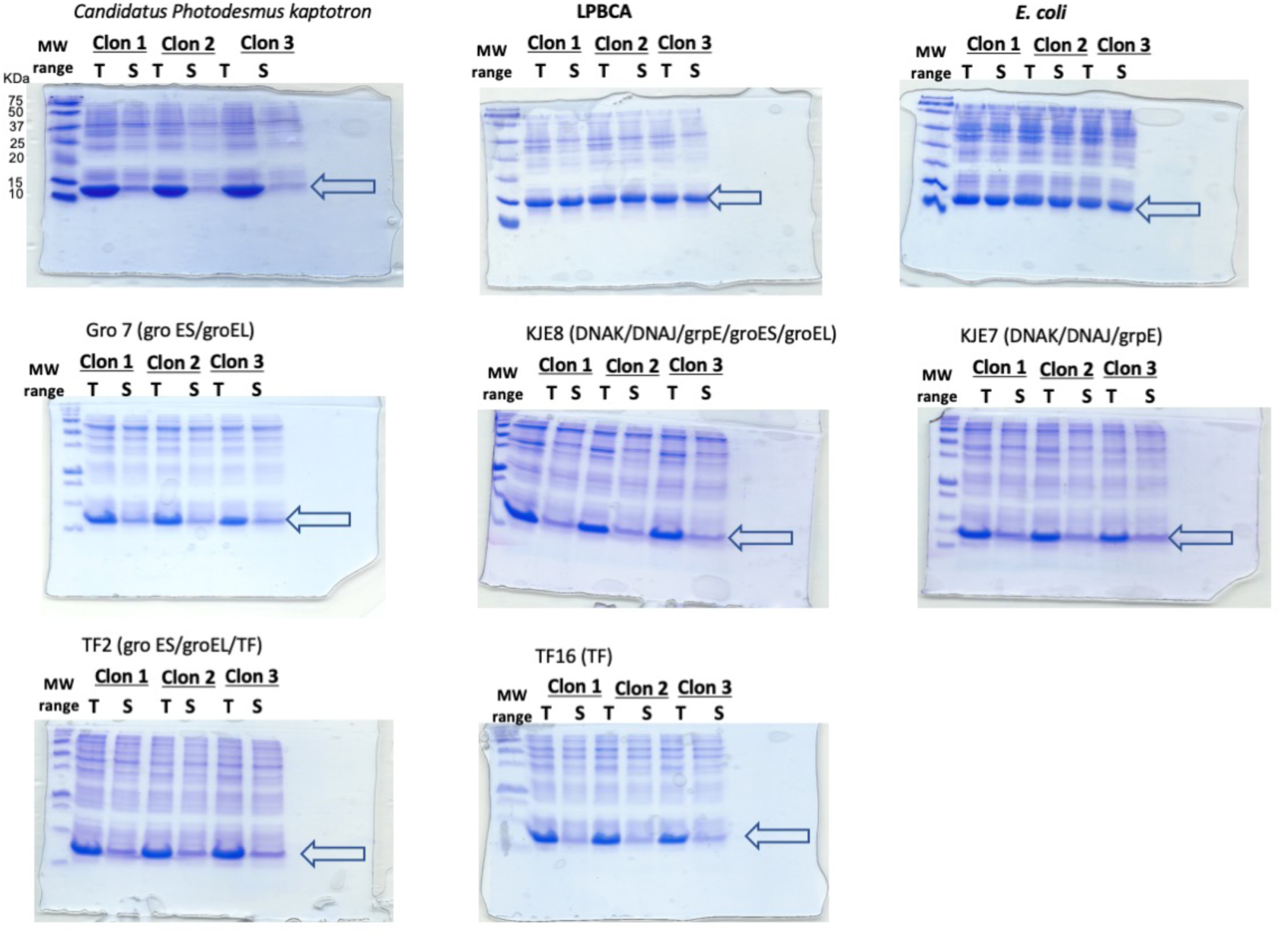
Electrophoresis SDS gels used for the determination of the fraction of soluble protein for *CPk* thioredoxin, the ancestral LPBCA thioredoxin and *E. coli* thioredoxin. For *CPk* thioredoxin, gels corresponding to experiments performed with over-expression of chaperone teams are also shown. Three independent determinations were carried out for each protein. T and S represent “total” and “soluble”. The molecular weights of the markers are shown in the gel for *CPk* thioredoxin.

**Figure S11.**
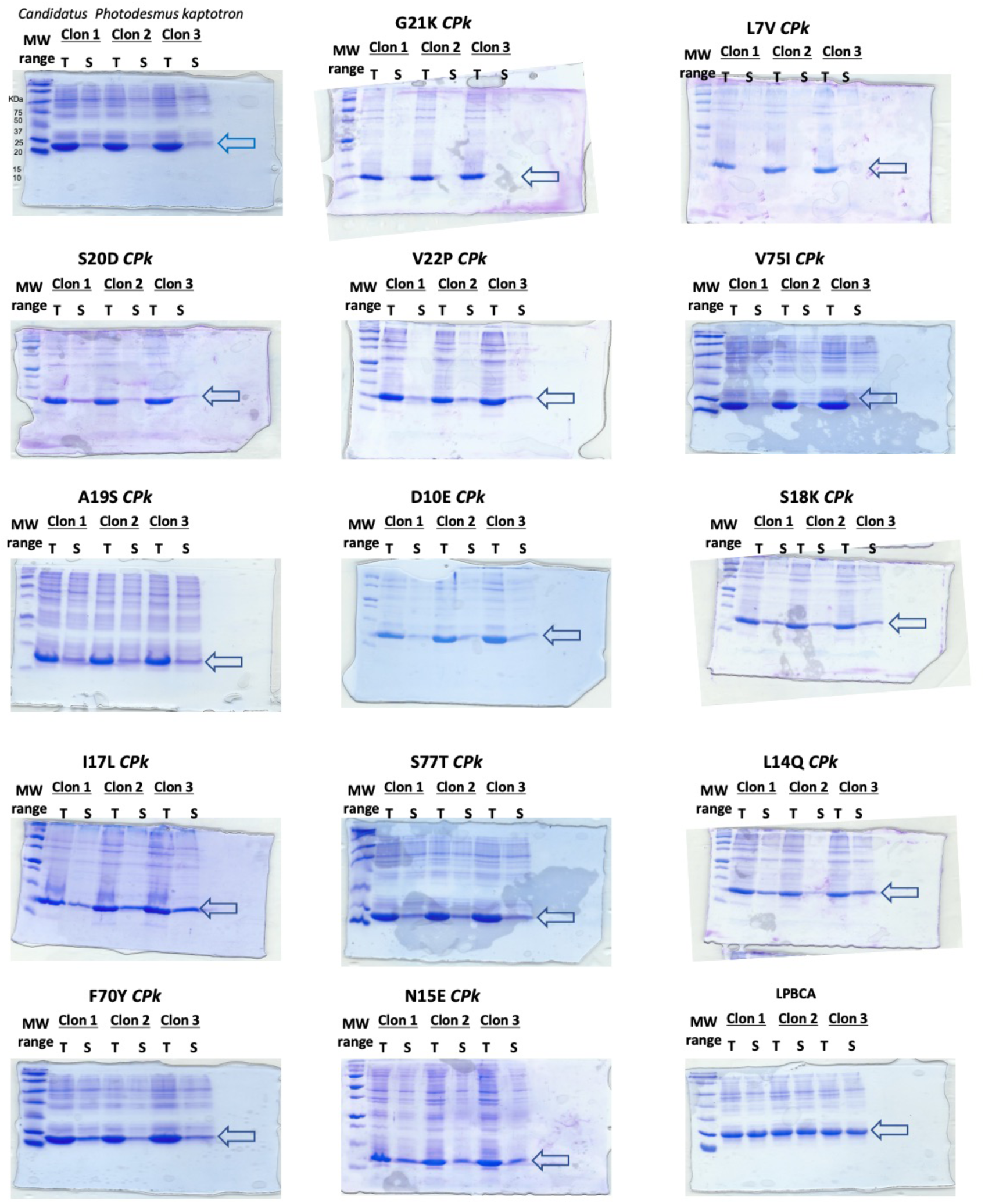
Electrophoresis SDS gels used for the determination of the fraction of soluble protein for *CPk* thioredoxin, single-mutant variants of *CPk* thioredoxin and the ancestral LPBCA thioredoxin. Three independent determinations were carried out for each protein. T and S represent “total” and “soluble”. The molecular weights of the markers are shown in the gel for *CPk* thioredoxin.

**Figure S12.**
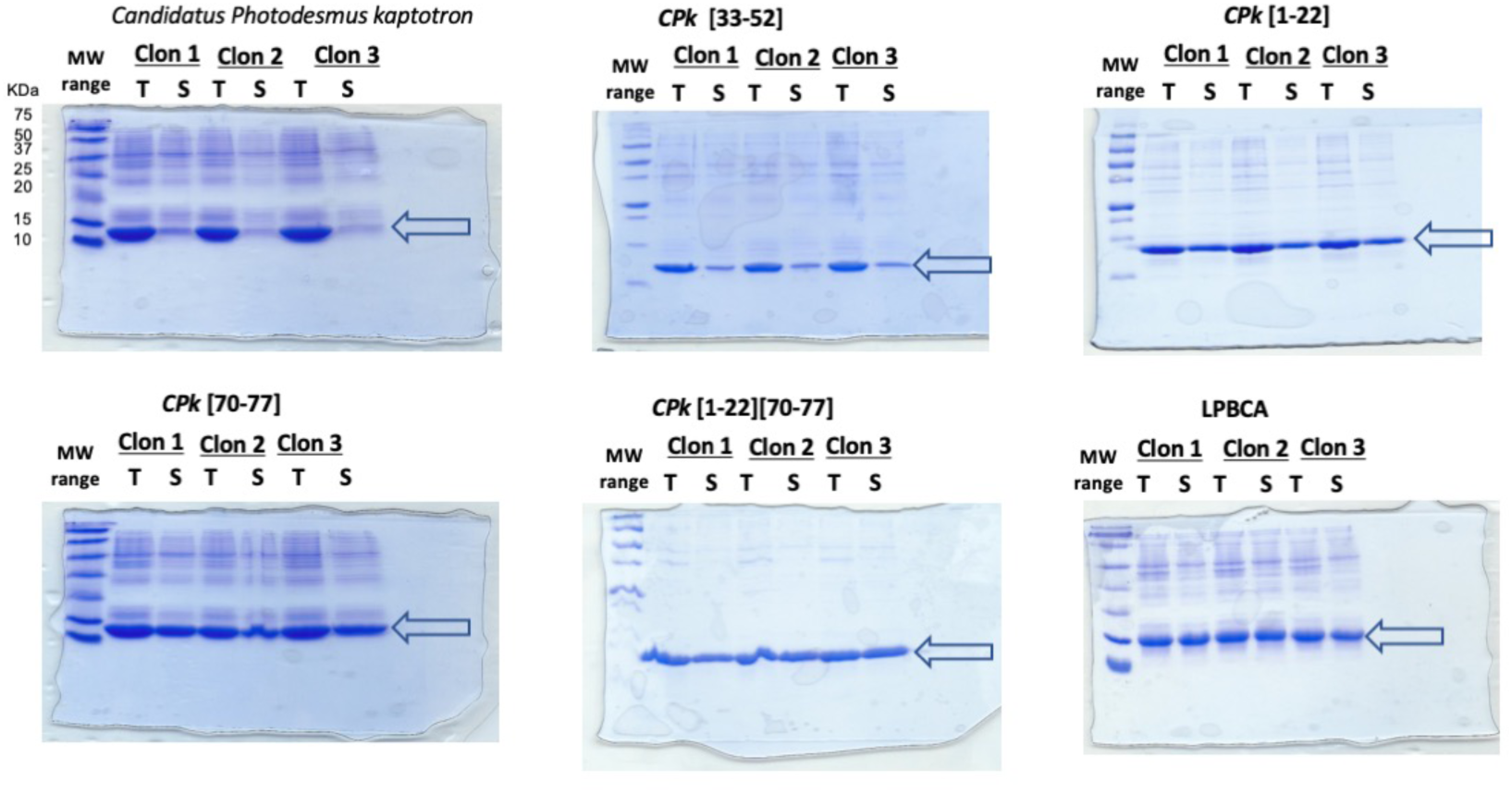
Electrophoresis SDS gels used for the determination of the fraction of soluble protein for *CPk* thioredoxin, the ancestral LPBCA thioredoxin and several modern/ancestral chimeras. Three independent determinations were carried out for each protein. T and S represent “total” and “soluble”. The molecular weights of the markers are shown in the gel for *CPk* thioredoxin.

**Figure S13.**
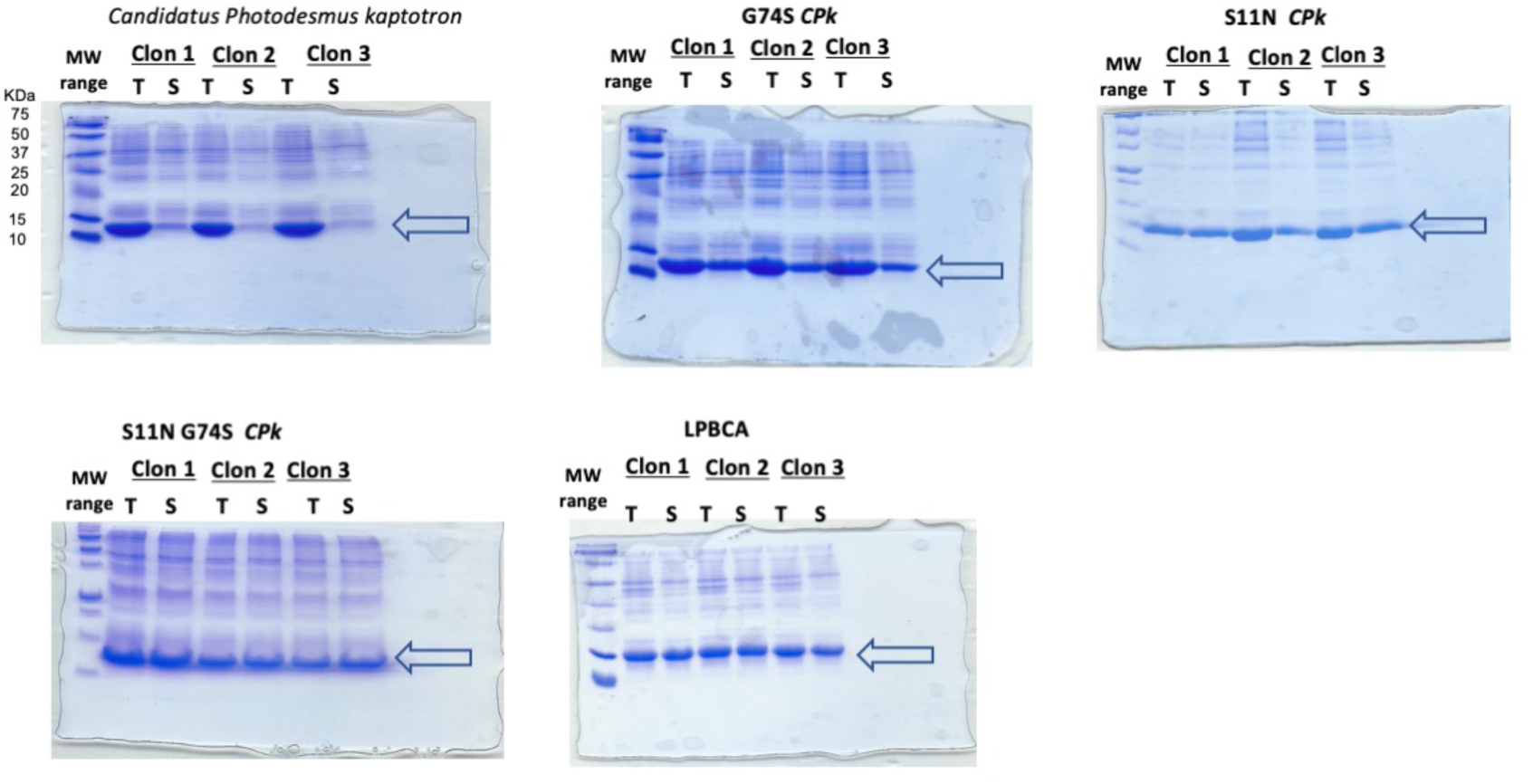
Electrophoresis SDS gels used for the determination of the fraction of soluble protein for *CPk* thioredoxin and its variants with single S11N and G74S mutations, as well as the double S11N/G74S variant of *CPk* thioredoxin. The ancestral LPBCA thioredoxin is also included for comparison. Three independent determinations were carried out for each protein. T and S represent “total” and “soluble”. The molecular weights of the markers are shown in the gel for *CPk* thioredoxin.

**Figure S14.**
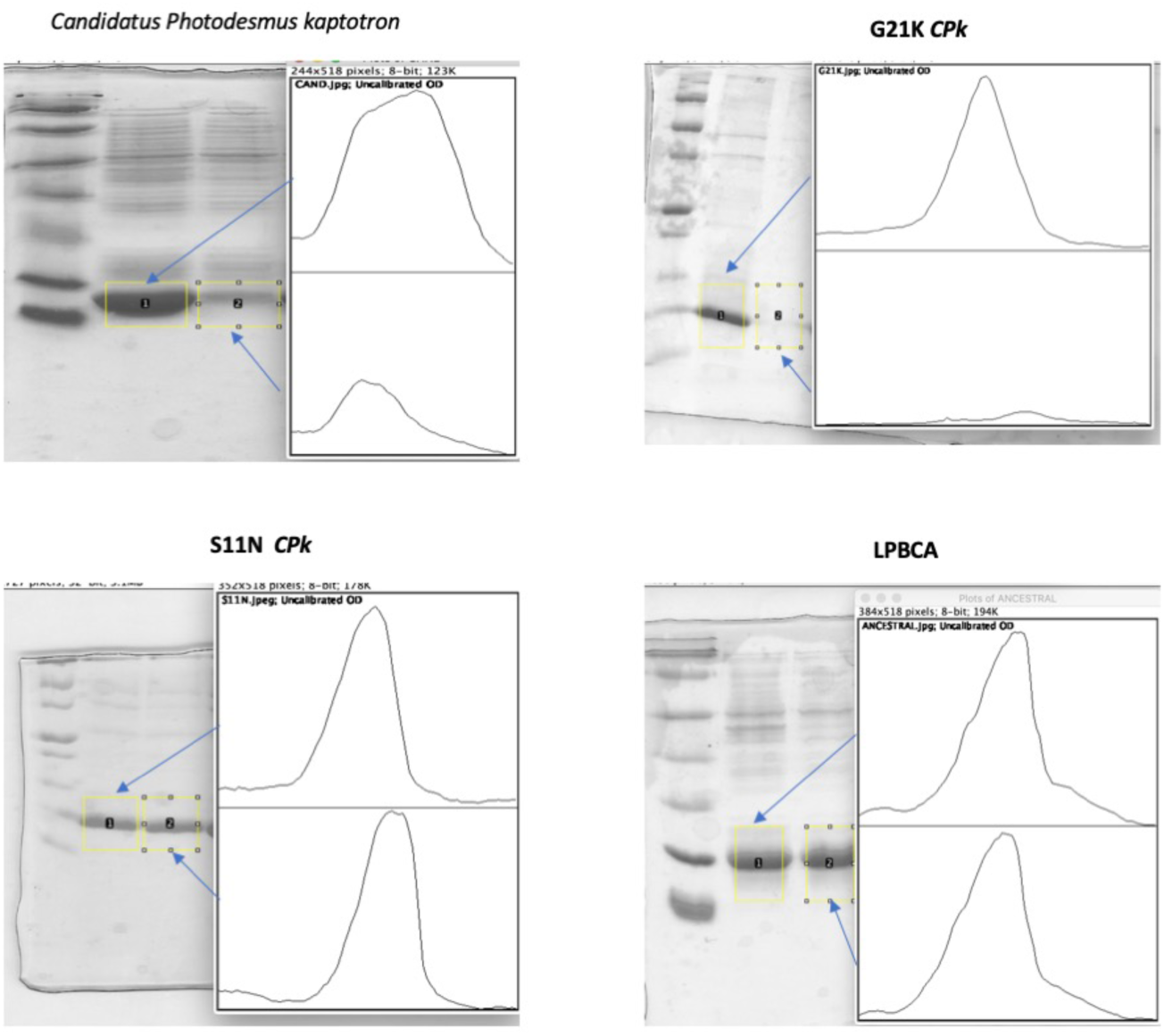
Representative examples of densitometric determination of amount of total and soluble protein from the SDS gels.

**Figure S15.**
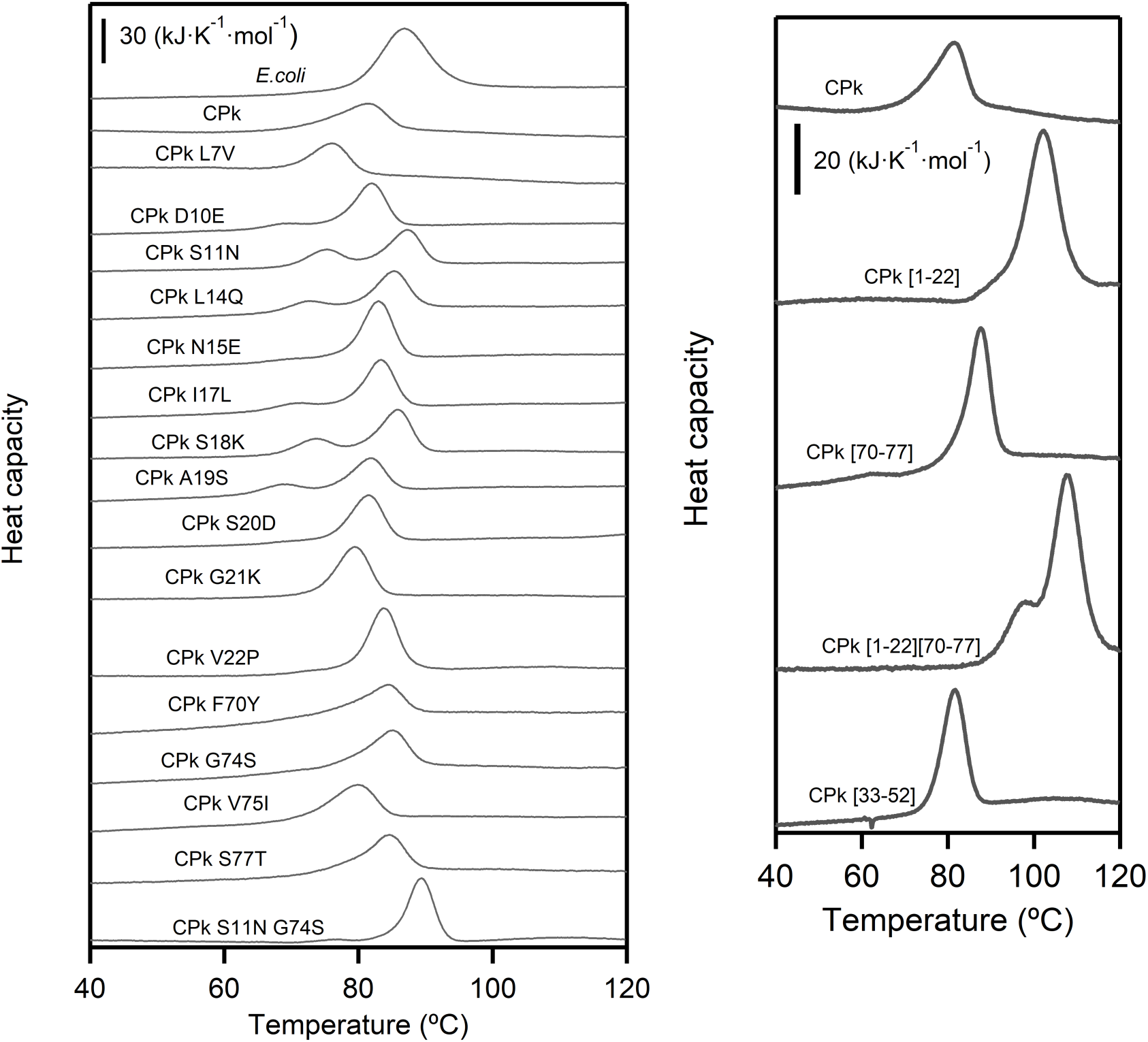
Differential scanning calorimetry profiles for the thermal denaturation of the thioredoxins studied in this work. Profiles have been shifted in the y-axis for representation purposes. Note that DSC instruments apply overpressure, which raises the boiling point of water and allows the temperature scan to extend above 100 °C. For a few variants, the calorimetric profiles show two peaks. This was found to be reproducible. In these cases, the temperature of the maximum of the major transition was used as a metric of stability.

